# TRPV1-lineage somatosensory fibers communicate with taste neurons in the mouse parabrachial nucleus

**DOI:** 10.1101/2021.10.17.464590

**Authors:** Jinrong Li, Md Sams Sazzad Ali, Christian H. Lemon

**Author notes:** The authors declare no competing interests, financial or otherwise. Portions of these data were presented in abstract form at the 2017 meeting of the Society for Neuroscience and the 2018 meeting of the Association for Chemoreception Sciences.

## Abstract

Trigeminal neurons supply somatosensation to craniofacial tissues. In mouse brain, ascending projections from medullary trigeminal neurons arrive at taste neurons in the autonomic parabrachial nucleus, suggesting taste neurons participate in somatosensory processing. However, the genetic cell types that support this convergence were undefined. Using Cre-directed optogenetics and *in vivo* neurophysiology in anesthetized mice of both sexes, here we studied whether TRPV1-lineage nociceptive and thermosensory fibers are primary neurons that drive trigeminal circuits reaching parabrachial taste cells. We monitored spiking activity in individual parabrachial neurons during photoexcitation of the terminals of TRPV1-lineage fibers that arrived at the dorsal spinal trigeminal nucleus pars caudalis, which relays orofacial somatosensory messages to the parabrachial area. Parabrachial neural responses to oral delivery of taste, chemesthetic, and thermal stimuli were also recorded. We found that optical excitation of TRPV1-lineage fibers frequently stimulated traditionally defined taste neurons in lateral parabrachial nuclei. The tuning of neurons across diverse tastes associated with their sensitivity to excitation of TRPV1-lineage fibers, which only sparingly engaged neurons oriented to preferred tastes like sucrose. Moreover, neurons that responded to photostimulation of TRPV1-lineage afferents showed strong responses to temperature including noxious heat, which predominantly excited parabrachial bitter taste cells. Multivariate analyses revealed the parabrachial confluence of TRPV1-lineage signals with taste captured sensory valence information shared across aversive gustatory, nociceptive, and thermal stimuli. Our results reveal that trigeminal fibers with defined roles in thermosensation and pain communicate with parabrachial taste neurons.

This multisensory convergence supports dependencies between gustatory and somatosensory hedonic representations in the brain.

## Introduction

Sensory systems organize the perceived world. Responses by sensory neurons order stimuli by qualitative features for discriminative sensory function. Sensory neurons also segment stimuli for autonomic recognition of their survival value. Examples of this include the innate preference for sweet taste tied to nutrient-rich sugars (Li et al., 2020b) and the aversiveness of pain or pain-related phenomena that may be associated with harmful stimuli (Walters and Williams, 2019).

Aversive sensations from multiple pathways engage cells in the parabrachial (PB) nucleus of the pons. In rodents, individual neurons in the lateral PB area respond to nociceptive stimuli applied to craniofacial regions and the limbs, respectively supplied by trigeminal and dorsal root afferents (Bernard and Besson, 1990). Lateral PB cells also respond to itch (Campos et al., 2018; Li et al., 2021), noxious visceral stimulation (Bernard et al., 1994; Campos et al., 2018), and auditory cues conditioned against fear (Campos et al., 2018). This spatial and multimodal convergence onto PB cells was discussed to convey an aversive valence common across sensations to engage protective behavioral and autonomic responses (Bernard and Besson, 1990; Gauriau and Bernard, 2002; Campos et al., 2018; Palmiter, 2018).

We recently found that the lateral PB area in mouse brain contains taste neurons that support a merger of input from gustatory and trigeminal circuits. A subpopulation of these PB cells responds to oral presence of behaviorally-avoided bitter taste stimuli and non-cell-type-selective excitation of the dorsal spinal trigeminal nucleus pars caudalis (Vc; Li and Lemon, 2019). Neurons in the dorsal Vc contribute to the trigeminoparabrachial tract (Cechetto et al., 1985) and respond to oral thermal and nociceptive stimuli (Carstens et al., 1998; Lemon et al., 2016). Along this line, lateral PB bitter taste neurons show responses to noxious oral heating that are reversibly suppressed by photoinhibition of presynaptic Vc circuits (Li and Lemon, 2019). Notably, Vc photoinhibition does not reliably affect PB neural responses to bitter tastes (Li and Lemon, 2019). This agrees with data showing Vc neurons are excited by oral thermal and nociceptive stimuli but not taste sensations (Simons et al., 2003b; Lemon et al., 2016), which engage PB cells through projections from the rostral nucleus of the solitary tract.

These results suggest that PB taste neurons that receive trigeminal projections represent sensory valence common to aversive gustatory (bitter) and oral thermal (noxious heat) sensations (Li and Lemon, 2019; Lemon, 2021). This convergence of cross-modal information onto common cells agrees with a role for the PB nucleus in protective coding. However, the genetic and functional definitions of trigeminal neurons that contribute to taste-somatosensory integrative circuits were undefined.

Here, we used Cre-directed optogenetics and *in vivo* neural recordings in mice to study whether a defined genetic type of thermosensory and nociceptive fiber contributes to trigeminal circuits that project to PB taste cells. We focused on trigeminal fibers marked by the capsaicin receptor: the transient receptor potential (TRP) ion channel TRP vanilloid 1 (TRPV1). Optogenetic stimulation of TRPV1-lineage afferents engages primary nociceptive neurons involved with protective thermosensory and nocifensive behaviors (Cavanaugh et al., 2011a; Mishra et al., 2011; Stemkowski et al., 2016; Browne et al., 2017; Black et al., 2020). We hypothesized that TRPV1-lineage fibers communicate with taste-active PB cells through a synaptic relay in the dorsal Vc, which projects to PB gustatory neurons (Li and Lemon, 2019). The Vc receives orofacial sensory input from afferents coursing the spinal trigeminal tract (Capra and Dessem, 1992), which is populated by TRPV1-lineage fibers (Cavanaugh et al., 2011a; Mishra et al., 2011).

We discovered that in the lateral PB area, traditionally defined taste neurons frequently respond to excitation of the central terminals of TRPV1-lineage fibers that arrive at the Vc. TRPV1-lineage afferents communicated with PB taste cells in a manner associated with gustatory and oral thermal tuning. We also uncovered evidence that the confluence of somatosensation with taste in PB circuits orders thermal and gustatory signals within a neural code for multisensory hedonic value.

## Materials and Methods

Overall, our approach was geared to understand if sensory messages from trigeminal TRPV1-lineage fibers are relayed by the Vc to PB taste neurons. Briefly, we recorded spikes from PB neurons in anesthetized mice and used Cre-directed optogenetics to excite the terminals of TRPV1-lineage primary neurons that arrived at the dorsal Vc (see Figure 1A). TRPV1 afferent terminals were predicted to stimulate Vc cells that project to the PB area. Responses to diverse taste and oral temperature stimuli were also recorded from PB neurons. Response data were, in part, compared between PB neurons that did or did not respond to optical excitation of TRPV1-lineage afferents. Additionally, taste preferences were measured in behaving mice to confirm the hedonic valence of select gustatory stimuli. Immunohistochemical and other methods validated mouse models.

**Figure 1.**
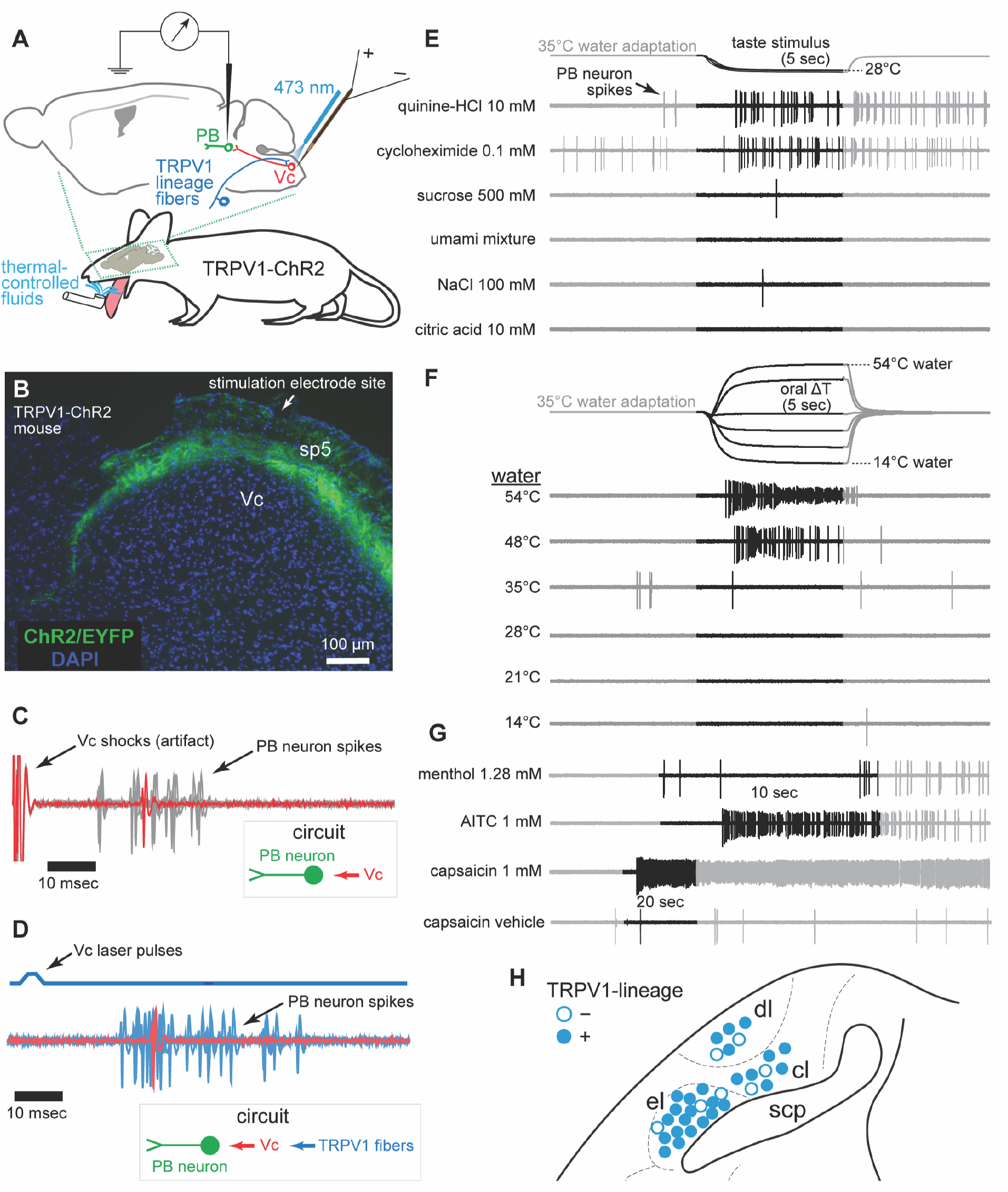
TRPV1-lineage afferents excite PB taste neurons. A) Drawing of experimental setup for PB neural recording and stimulation of TRPV1-lineage afferents that contacted Vc neurons projecting to PB cells. Vc electrical stimulation and delivery of oral fluids are also depicted. B) Coronal brain section shows EYFP labeling of TRPV1-lineage fibers entering the superficial dorsal Vc from the spinal trigeminal tract (sp5). C) Spikes recorded from a single PB neuron to 1 Hz electrical pulse stimulation of the dorsal Vc. Multiple sweeps are superimposed, with an example in red. Inset shows the neural circuit predicted to underly the spikes. D) Spikes recorded from the same neuron to 1 Hz laser pulse stimulation directed to tissues above the dorsal Vc, which engaged arriving TRPV1-lineage afferents. This neuron was TRPV1-lineage positive. E) Responses by the neuron in C and D to taste solutions at 28°C. F) Responses by the same neuron to change in water temperature flowed into the mouth. G) Responses by the neuron in C-F to oral delivery of chemesthetic and chemonociceptive stimuli. H) Coronal view of recording locations for PB neurons (circles) where sites were recovered. PB areas: el, external lateral; dl, dorsal lateral; cl, central lateral; scp, superior cerebellar peduncle. Rostrocaudal dimension is conflated for simplification.

### Mice

For neurophysiological studies, we used Cre-lox technology to generate TRPV1-ChR2 mice, which express channelrhodopsin-2 (ChR2) and EYFP in all TRPV1-lineage fibers. The present study focused on TRPV1-lineage fibers coursing the trigeminal tract. TRPV1-ChR2 mice support selective excitation of TRPV1-lineage primary neurons via blue light activation of ChR2 and are well-established in studies of nociception (Stemkowski et al., 2016; Browne et al., 2017; Black et al., 2020). TRPV1-ChR2 mice were the F1 progeny of a cross between homozygous TRPV1^Cre^ mice, which express Cre recombinase in TRPV1-lineage neurons (JAX #017769; Cavanaugh et al., 2011b), and homozygous Ai32 mice, which direct Cre-dependent transcription of a ChR2/EYFP fusion gene (JAX #024109; Madisen et al., 2012). Our PB neural recordings used 70 adult TRPV1-ChR2 mice, including 41 females (body weight [mean ± SD]: 24.6 ± 2.3 grams) and 29 males (33.7 ± 5.1 grams). Additional Ai32, TRPV1^Cre^, and also mT/mG fluorescent Cre-reporter mice (JAX #007676) were used in procedural control studies. For behavioral studies, 6 adult C57BL/6J mice (JAX #000664, 3 females [22.2 to 23.5 grams, at study onset] and 3 males [27.9 to 30.1 grams]) participated in brief-access fluid licking tests with taste solutions.

All procedures performed on mice were approved by the University of Oklahoma Institutional Animal Care and Use Committee and followed National Institutes of Health guidelines. Mice entered studies as naïve to experiments and were housed in a vivarium that maintained a 12:12 hour light:dark cycle and an air temperature of ∼20°C. Food and water were available ad libitum except during water restriction conditions in behavioral tests.

### Immunohistochemistry

We performed GFP/YFP immunohistochemistry on medullary tissue from Cre-negative Ai32 mice to confirm there was no off-target or leak (i.e., Cre-independent) expression of ChR2/EYFP (Madisen et al., 2012). As a control, we applied the same immunohistochemical procedures to Cre-positive tissues from TRPV1^Cre^;mT/mG mice. TRPV1^Cre^;mT/mG mice were generated by crossing TRPV1^Cre^ mice with the mT/mG fluorescent Cre-reporter line. TRPV1^Cre^;mT/mG offspring express EGFP in TRPV1-lineage primary neurons and axons.

Mice used in immunohistochemistry studies were overdosed with sodium pentobarbital (270 mg/kg, i.p.). Mice then received transcardial perfusion of filtered 0.9% NaCl (10 mL) followed by 4% (w/v) paraformaldehyde and 3% (w/v) sucrose in 100 mM phosphate buffer (80 mL). Brains were removed and stored in this fixative for ∼4 hours. Brains were then transferred to 30% (w/v) sucrose in 100 mM phosphate buffer and kept at 4°C until sectioning.

Serial coronal sections (20 µm) of brain tissue were cut on a microtome (Leica SM2010R). Brain sections were washed three times in 10 mM phosphate-buffered saline with 0.05% Tween-20 (PBST) for 5 min and incubated in permeabilization solution (0.25% NP-40) in PBST for 10-20 minutes. Sections were then triple washed with PBST and incubated in a blocking solution (5% normal goat serum in PBST) for 2 hours at room temperature. Next, sections were again triple washed with PBST and then incubated overnight at 4°C with a primary rabbit anti-GFP/YFP antibody (1:1000, Abcam, ab6556). After this, sections were washed five times with PBST and then incubated for 1 hour in PBST with a secondary antibody: Alexa Fluor® 647 goat anti-rabbit (1:2000, Invitrogen, A-21244). Approximately one-half hour later, DAPI (3 µg/ml) was added to the secondary incubation wells. Finally, brain tissue was rinsed at least three times in 100 mM PBS for more than 10 min. Sections were mounted onto clean glass slides, air dried, and cover-slipped with a mixture of 80% (v/v) glycerin and 2.5% (w/v) triethylenediamine (anti-fading agent, Sigma-Aldrich, D27802) in 10 mM PBS. Fluorescent images were acquired using a compound microscope (Zeiss Apotome).

### Neurophysiology preparation

TRPV1-ChR2 mice were prepared for stimulation of the Vc and PB neural recording following initial sedation with isoflurane in O_2_ and then induction with ketamine (100 mg/kg, i.p.) and xylazine (10 mg/kg, i.p.). Atropine (24 µg/kg, i.p.) was administered to reduce bronchial secretions. A tracheostomy tube (PE 60) was inserted to allow breathing during oral stimulation with liquids (Li et al., 2020a). A small suture was placed through the superficial ventrorostral tongue and the mandibular incisors trimmed for tongue extension. Mice were secured in a stereotaxic instrument with ear bars (Model 930, David Kopf Instruments, Tujunga, CA). To maintain anesthesia during experiments, mice freely inhaled 0.8 to 1.2% isoflurane in O_2_ through their tracheostomy tube using a custom device (Li et al., 2020a). A feedback-controlled heating pad kept body temperature at ∼37°C. Blood oxygenation and heart rate were monitored using a pulse oximeter.

Under anesthesia, a midline incision was made on the scalp to expose bregma and lambda, which were brought into the same dorsoventral plane. The mandible was gently deflected downward, and the tongue was extended from the mouth by light pressure on the ventrorostral suture. Using stereotaxic coordinates, a small unilateral craniotomy was made on the skull to allow dorsoventral electrode access to the PB area. The ipsilateral caudal zone of the occipital bone was trimmed to access the Vc.

A custom optrode was used to optically excite the terminals of TRPV1-lineage afferents that arrived at the at the rostrocaudal level of the orosensory dorsal Vc. The optrode had a fiber optic and electrode channel, with the latter supporting measurement of coarse neural activity and weak electrical stimulation of brain tissue. The optrode was constructed by pairing a concentric bipolar electrode (CEA-200, Microprobes, Gaithersburg, MD) with the prepared end of a 200 μm diameter fiber optic cable (0.39 numerical aperture, ThorLabs, Newton, NJ). The tip of the fiber optic probe was recessed relative to the electrode tip.

The optrode was targeted to the orosensory Vc using both coordinates and electrophysiological guidance. Using a micromanipulator (SM-15R, Narishige International, Amityville, NY), the optrode approached the Vc at ∼115° sagittal to accommodate electrode access to the PB area. Vc coordinates were from our prior work: ∼8.0 mm caudal of bregma and ∼1.8 mm lateral of the midline (Lemon et al., 2016; Li and Lemon, 2019). The cerebellum was not disturbed to target the Vc. The electrode tip of the optrode was located at a superficial depth in the dorsal Vc (see Figure 1B) where orosensory neurons reside (Carstens et al., 1998; Lemon et al., 2016). The recessed fiber optic probe of the optrode was then able to deliver laser light pulses that penetrated the medullary surface near the electrode tip (Li and Lemon, 2019). Just below this surface is the spinal trigeminal tract, which carries TRPV1-lineage afferents (Cavanaugh et al., 2011a; Mishra et al., 2011). Final positioning of the optrode electrode in the orosensory Vc was confirmed by monitoring for multiunit responses to oral presentation of cool water following 35°C adaptation, as below.

Following optrode placement, a single channel tungsten recording electrode (2 to 5 MΩ, FHC Inc., Bowdoinham, ME) was located in the PB area at ∼90° sagittal/lateral using stereotaxic coordinates. We targeted lateral PB regions where the confluence of taste and trigeminal somatosensory pathways arises (Li and Lemon, 2019). Coordinates were 4.7 to 5.1 mm caudal of bregma, 1.1 to 1.4 mm lateral of the midline, and 2.2 to 3.0 mm below the brain surface. The electrode was advanced dorsoventrally using an electronic micro-positioner (Model 2660, David Kopf Instruments). We monitored neural activity through this electrode and sought out PB taste neurons by measuring responses to oral delivery of a room temperature aqueous solution of 300 mM NaCl. This stimulus excites diverse groups of mouse PB gustatory neurons oriented to electrolyte, aversive bitter, or appetitive taste stimuli (Li and Lemon, 2019).

Electrophysiological activity recorded through the high-impedance PB electrode was AC amplified (P511 with high-impedance probe, Grass Technologies) and band-passed to ∼0.3 to 10 kHz. A template-matching algorithm sampled extracellular spikes from well-isolated single neurons at 25 kHz (1401 interface and Spike2 software version 9, CED, Cambridge, England). Spikes were time stamped to 0.1 msec. Data files were stored and analyses performed offline.

### Optical and electrical stimuli

Individual PB neurons were tested to respond to blue laser (473 ± 1 nm DPSS laser, OEM Laser Systems, Inc., East Lansing, MI) pulses (∼5 mW/4 msec; 1 Hz) delivered through the optrode to the medullary surface above the orosensory dorsal Vc. Laser pulse trains were intended to excite ChR2-positive TRPV1-lineage fibers that arrived at the orosensory Vc via the spinal trigeminal tract. The central terminals of TRPV1-lineage afferents were predicted to arborize (Contreras et al., 1982) with Vc cells that composed the trigeminoparabrachial circuit projecting to the PB area. The laser was controlled by programmable TTL coding from the data acquisition system. Downstream PB neurons that reliably responded to laser pulses, as below, were considered to receive input from Vc neurons that relayed sensory messages from TRPV1-lineage fibers to PB cells.

PB neurons were also tested to spike to weak electrical pulses (∼150 μA/500 μsec; 1 Hz) delivered through the optrode to the orosensory dorsal Vc. This supported comparison of evoked responses in PB cells between non-selective (electrical) and TRPV1-lineage specific (laser) stimulation of Vc circuits. Constant current pulses were generated by a programmable stimulator (S88X and PSIU6X, Grass Technologies). The appearance of electrically-evoked spikes, and offline statistical verification of their reliability as below, provided evidence that PB neurons received orthodromic input from upstream Vc projection circuits (Li and Lemon, 2019).

### Taste, temperature, and chemesthetic stimuli

Neurons were tested with oral delivery of taste, temperature, and chemesthetic stimuli in sequence over three separate blocks. Within each block, stimulus trials were randomly ordered, without replacement, for each neuron. The sole exception was capsaicin, which was always tested at the end of the chemesthetic stimulus array due to the lingering post-stimulus effects of this nociceptive agent on PB cells (Li and Lemon, 2019).

Taste stimuli were aqueous solutions of 500 mM sucrose, an umami mixture of 100 mM monopotassium glutamate and 10 mM inosine 5’-monophophate, 100 mM NaCl, 10 mM citric acid, 10 mM quinine-HCl, and 0.1 mM cycloheximide. These tastants are associated with the sweet (sucrose), umami, salty (NaCl), sour (citric acid), and bitter (quinine and cycloheximide) human taste categories. Taste chemicals were of high purity (Sigma-Aldrich, St. Louis, MO) and dissolved in purified water.

Concentrations of sucrose, NaCl, citric acid, and quinine parallel those used in prior neurophysiological studies of gustatory processing, including recordings carried out in different species or neural structures (e.g., Breza et al., 2006; McCaughey, 2007; Fletcher et al., 2017; Martin et al., 2021) and in the mouse PB area (Tokita and Boughter, 2016; Li and Lemon, 2019). The selected 0.1 mM concentration of cycloheximide causes a neural response comparable to 10 mM quinine in mouse medullary bitter taste neurons dually sensitive to these stimuli (Wilson et al., 2012). Notably, across-neuron patterns of response to cycloheximide or quinine remain correlated and distinct from non-bitter taste stimuli across a broad range of concentrations (Wilson et al., 2012). Finally, concentrations for the components of the umami mixture were selected to induce gustatory activity comparable to 500 mM sucrose in mouse PB neurons (Tokita and Boughter, 2016).

Taste solution temperature was controlled during neural data acquisition, with stimuli delivered at 28°C following oral adaptation to 35°C water, as measured below. Gustatory stimuli were delivered at a temperature less than adaptation temperature to reflect thermal shifts that may happen when sampling an external taste stimulus. This approach was used in our prior thermogustatory neurophysiology studies in mice (Li and Lemon, 2015; Lemon et al., 2016; Li and Lemon, 2019) and follows data from humans that imply cooling can accompany taste sensations in mammals due to the relatively high resting temperature (near 36°C) of the mouth (Green, 1986).

Our delivery method for tastes also gauged neural activity that may be due to the temperature instead of chemical component of taste solutions. Gustatory neural recording studies in rodents usually proffer taste fluids at uncontrolled temperatures less than 35°C, typically at a substantially lower laboratory room temperature. Notably, fluids presented orally to mice at temperatures <30°C stimulate primary trigeminal cooling fibers (Yarmolinsky et al., 2016; Leijon et al., 2019) and cooling-sensitive trigeminal neurons in the Vc (Lemon et al., 2016). This opens the possibility that PB gustatory neurons receiving Vc input may respond to thermal features of taste solutions. To account for this and measure chemical taste activity, we recorded PB neural responses to taste stimuli delivered at a controlled temperature of 28°C. On separate trials, we measured activity to the 28°C water vehicle, with gustatory firing corrected for potential activity to the vehicle as below. Temperature-corrected and -uncorrected taste responses were also compared, defined below.

For temperature-only trials, thermal stimulation was achieved using oral flow of purified water at 14°, 21°, 28°, 35°, 48°, and 54°C following oral adaptation to 35°C water. All fluid temperatures were measured at the moment of oral entry using a miniature thermocouple probe (IT-1E, time constant = 0.005 sec, Physitemp Instruments, Inc., Clifton, NJ) and thermometer (BAT-12, precision = 0.1°C, Physitemp Instruments, Inc.). Temperatures below 35° were considered cool whereas 48° and 54°C exceed noxious heat threshold and engage nociceptors (Caterina, 2007). Temperature data were sampled (1 kHz) by the data acquisition system in synchronicity with neural spike trains (see Figure 1E, 1F). Each stated temperature is the temperature that fluid entering the mouth reached during the stimulus period, averaged across trials for all neurons.

Chemesthetic stimuli (Sigma-Aldrich) were 1.28 mM 28°C (–)-menthol, 0.1 mM 35°C allyl isothiocyanate (AITC; also known as mustard oil), 1 mM 35°C AITC, and 1 mM capsaicin at room temperature. These stimuli/concentrations induce varied orosensory behavioral reactions in mice and were tested to further explore PB hedonic processing.

Menthol is an agonist of the cold receptor TRP melastatin 8 (TRPM8) (McKemy et al., 2002). The tested concentration of menthol stimulates lingual trigeminal fibers (Lundy and Contreras, 1995) and its oral sensation is not avoided by mice. In an orosensory setting, wild-type mice show only a mild reduction in licking 1.28 mM menthol compared to water (Lemon et al., 2019).

AITC engages nociceptors expressing TRP ankyrin 1 (TRPA1) and stimulates TRPV1 at high concentrations (Jordt et al., 2004; Everaerts et al., 2011). Whereas 0.1 mM AITC causes a moderate reduction in licking compared to water in mouse orosensory tests, 1 mM AITC elicits near-zero licks and strong aversion (Lemon et al., 2019).

Capsaicin is a selective agonist of the heat-sensitive nocisensor TRPV1 (Caterina et al., 1997). The selected 1 mM concentration of capsaicin follows prior electrophysiological studies of capsaicin effects on rodent lingual trigeminal neurons (Carstens et al., 1998) and causes strong orosensory aversion in mice (Ellingson et al., 2009; Long et al., 2010).

Concentrations of capsaicin and the other chemesthetic agents may be reduced at receptor sites due to diffusion through oral epithelia (Simons et al., 2003a). All chemesthetic stimuli were dissolved in purified water except for capsaicin, which required a vehicle solution of 1.5% ethanol/1.5% Tween 80 in purified water. Neurons were also tested with the vehicle-only (without capsaicin) solution on separate trials.

Aside from capsaicin, stimulus fluids were delivered to oral epithelia using a custom flow apparatus that adapted the mouse oral cavity to 35°C water and then switched solution flow to temperature-controlled stimuli, as described (Wilson and Lemon, 2014; Lemon et al., 2016). Oral flow rates for fluids were ∼1.5 mL/sec. Prior to testing, all stimulus solutions were kept in airtight glass bottles placed in programmable heating and cooling water baths to control temperature. Temperature changes in the mouth were rapid and stable during stimulus periods (see Figure 1E, 1F). Taste and temperature stimuli were presented for 5 sec. Menthol and AITC solutions were presented for 10 sec. Trials ended 5 sec after the stimulus period. Fifteen neurons were tested with 20 sec presentations of menthol and AITC at 35°C on longer trials, albeit no interpretative differences emerged. Stimuli bathed anterior and posterior oral fields (i.e., whole-mouth stimulation; Wilson et al., 2012). The adaptation rinse resumed following the stimulus period and continued during inter-trial intervals (∼2 min) to facilitate thermal adaptation of oral tissue to physiological levels. Mice did not ingest adaptation or stimulus fluids, which entered the mouth and then fell into a drain beneath the mandible.

On capsaicin and capsaicin vehicle-only trials, the adaptation rinse was paused and a bolus of capsaicin or the vehicle solution was brushed onto the dorsorostral tongue for 20 sec using a disposable cotton-tipped applicator (Fisherbrand, 22-363-157). Prior to testing, the cotton bulb of the applicator was stretched/puffed by the experimenter to be approximately 1.5 cm long and 0.5 cm wide at the tip. This facilitated absorption of capsaicin or the vehicle solution and acted a soft “brush” for chemical delivery to the tongue. A new applicator was used for each trial, with 150 μL of capsaicin or the vehicle solution applied to the cotton tip immediately before use. Following lingual stimulation, the 35°C adaptation rinse resumed and neural activity was monitored for an additional 80-95 sec.

The capsaicin vehicle-only trials were used to capture extraneous neural activity that arose during the capsaicin delivery process. This process includes light lingual brushing (mechanical stimulation) and tongue cooling, as pausing the 35°C adaptation rinse exposed the warmed tongue to cooler room temperature air while it extended from the mouth. As below, neural responses on capsaicin trials were corrected for activity to mechanical brushing or cooling by subtracting spikes on vehicle-only trials from the evoked capsaicin response. To quantify neural firing on vehicle-only trials, activity was corrected using an additional control trial, referred to as a blank trial, where the warm adaptation rinse was paused for 20 sec but no lingual stimulus was applied. This trial accounted for spikes attributable to tongue cooling.

### Parabrachial histology

After data were collected from the last PB neuron of the day, mice were overdosed with sodium pentobarbital (270 mg/kg, i.p.) and weak current (100 μA/1.5 sec) was passed through the recording electrode tip to mark its final position. Brains were fixed by transcardial perfusion using filtered isotonic saline followed by 4% paraformaldehyde/3% sucrose. Brains were removed and stored in a 4% paraformaldehyde/20% sucrose solution. Coronal sections (40 μm) from each brain were cut using our microtome, mounted onto slides, and stained for histological analysis of electrode placement using anatomical landmarks (Franklin and Paxinos, 2008). Many, but not all, recording sites were marked or recoverable.

### Brief-access taste preference tests

Water-restricted C57BL/6J mice completed brief-access fluid licking tests for select taste solutions in a Davis Rig contact lickometer (Med Associates Inc., Fairfax, VT). Brief-access tests gauge initial licking responses to small volumes of stimulus fluid and measure how oral sensations influence ingestive preference behaviors (Smith, 2001). The lickometer afforded measurement of initial licking responses to multiple concentrations of one taste stimulus within each daily test session, which lasted ∼20 minutes. Taste stimuli were room temperature aqueous solutions of citric acid (0 [water vehicle], 1, 3, 10, 30, and 100 mM), NaCl (0, 10, 30, 100, 300, 500, 1000 mM), and sucrose (0, 10, 100, 300, 500, 1000 mM).

For all brief-access tests, we followed water restriction, training, and testing procedures generally described in Lemon et al. (2019). Briefly, mice were individually housed and subjected to water restriction conditions where they obtained all daily fluids in the lickometer. Food was always available in their home cage. All mice maintained approximately 80% or more of their baseline body weight during testing. Daily test sessions allowed 10 sec access to each tastant concentration, proffered 3 times across randomized blocks of trials of the test solution. For each mouse, software coupled to the lickometer recorded the number of licks they made to the fluid presented on each trial. Mice were first tested with the concentration series of citric acid for 7 consecutive days. Next, mice underwent brief-access tests with the NaCl series, which was tested for 7 consecutive days. The sucrose concentration series was tested last and for 5 consecutive days. Mice were given at least 2 days of *ad lib* access to water in between taste stimulus tests.

## Experimental Design and statistical analysis

### Neural sample

A total of 106 PB neurons were recorded from TRPV1-ChR2 mice. Not every cell remained isolated across all stimulus conditions and some analyses involved subsets of neurons. Sample sizes for each analysis are reported. Considering 70 neurons tested with all stimuli, sex did not influence responses to temperature (non-significant sex × stimulus interaction: *F*_5,340_ = 1.1, *p* = 0.4), taste (*F*_5,340_ = 0.2, *p* = 0.8), or chemesthetic (*F*_3,204_ = 0.8, *p* = 0.4) stimuli (two-way ANOVAs with *p*-levels corrected by the Greenhouse-Geisser method for lack of sphericity). Sex was not further analyzed as a factor. For multiple neurons sequentially recorded from the same mouse, their responses to taste stimuli did not show positive correlation (Spearman’s rank-order correlation coefficient ≤ 0.29, *p* ≥ 0.23). These neurons were presumed to display statistical independence in their firing characteristics and, accordingly, were analyzed as independent units.

### PB neural responses to Vc electrical and laser pulses

For individual PB neurons, a statistical approach determined if laser and electrical pulses delivered to the orosensory dorsal Vc caused a significant response. This evaluated coupling of PB cells with upstream Vc circuits and TRPV1 afferents. Electrical and laser pulses were applied over many trials at 1 Hz, with 40 electrical and 50 laser pulses tested during separate trial blocks. PB neural responses to electrical and laser pulse trains were analyzed separately. Each pulse marked the beginning of a trial and started a clock at 0 msec to measure the time of occurrence of any spikes that followed. Spike times across trials were conflated into a vector. An iterative Poisson method determined if the evoked spike rate built from sequential spikes in this vector was significantly greater than expected by chance, as described (Chase and Young, 2007; Wilson and Lemon, 2014). Evoked response latency was defined as the time of the post-pulse spike where the firing rate became significant (Li and Lemon, 2019).

A PB neuron that significantly responded to Vc electrical pulses was considered to receive input from Vc-parabrachial projection circuits. A neuron that significantly responded to Vc laser pulses was evidenced to receive input from Vc neurons that relayed information from TRPV1-lineage fibers to PB cells. These PB neurons were classified as TRPV1-lineage positive. PB neurons that did not significantly respond to Vc laser pulses were classified as TRPV1-lineage negative. TRPV1-lineage negative PB neurons usually showed a clear lack of evoked spikes during recordings. Importantly, TRPV1-lineage negative neurons were included in analyses only if Vc laser stimulation excited other PB neurons in the same mouse. This established positive control for the experimental preparation and circuit analysis.

### Thermal and chemosensory responses in PB neurons

Responses by individual PB neurons to each oral temperature and chemosensory stimulus were initially quantified by counting spikes from stimulus onset to trial end. For temperature trials, responses to 14°, 21°, 28°, 48°, and 54°C water were then corrected for baseline activity by subtracting the number of spikes to 35°C water, which was the adaptation rinse. Responses to 28°C taste stimuli were corrected for activity to the cooling feature of taste solutions by subtracting the number of spikes to the taste stimulus vehicle, 28°C water. Likewise, responses to menthol and AITC solutions were corrected for vehicle activity by subtracting spikes to isothermal water. Responses to capsaicin were quantified as spikes that emerged during and after stimulus application minus spikes to the capsaicin vehicle. Finally, PB neural activity on the capsaicin vehicle-only trials was calculated as spikes that appeared during and after brushing the vehicle to the tongue minus spikes on the blank trial.

In one analysis, temperature-corrected and -uncorrected taste responses were compared to study if the thermal component of taste solutions produced a neural effect. For temperature-uncorrected taste activity, the response each neuron showed to the taste solution vehicle, 28°C water, was not subtracted from its measured taste responses. All other analyses used temperature-corrected taste activity.

All firing rates were quantified in spikes per sec (Hz). The breadth of neural tuning to taste stimuli was calculated using lifetime sparseness (Iurilli and Datta, 2017), which ranges from 0 (broadly responsive across stimuli) to 1 (selective firing). Heatmaps normalized responses for visualization by making the smallest response in each array the first shade of the colormap, and the largest response the last shade.

### Clustering neurons by orosensory responses

PB neurons were clustered by sensory tuning using an approach based on non-negative matrix factorization (NMF). NMF is a dimensionality reduction technique that can find a small number of additive features in a data set that combine to form its overall pattern (Lee and Seung, 1999; Brunet et al., 2004). In our case, these additive features were the responses to orosensory stimuli that tend to co-stimulate PB cells. Identification of these stimulus sets was used to cluster PB neurons based on similarities in sensory tuning. Most PB neurons showed excitatory sensory responses (>0 Hz), which agrees with the additivity constraint of data components for NMF.

Our use of NMF for neuron classification largely followed the study of Brunet et al. (2004), which used NMF in a different yet related setting. In brief, we applied NMF to a stimulus × neuron response matrix, where responses less than 0 Hz were converted to zero. From this matrix, NMF generated two non-negative, lower-dimensional matrices parameterized in part by *k*, which is the expected number of clusters present in the neuronal sample. The low-dimensional matrices partly weighted how well PB neurons fit with each cluster, which reflected a commonality in sensory responsiveness for included cells. Neurons were assigned to the cluster where their weighted fit was the highest. NMF was performed using the *nnmf* function in MATLAB with default settings.

The number of cell clusters present in the data was found by first repeatedly computing NMF from random starting configurations across different values of *k*. We used *k* = 2 to 5 for classification of neurons by responses to tastants, and *k* = 2 to 7 for classification based on responses to gustatory and thermal stimuli. NMF was ran 100 times for each *k*. Infrequently, some runs converged to a rank lower than *k* and were discarded. To select final clusters, we used the highest *k* that resulted in reliable classification of cells across successful NMF runs, as validated using visual and quantitative methods (Brunet et al., 2004). Briefly, consensus matrices used entries with shades of blue to show the percentage of runs that placed pairs of neurons into the same cluster. Deep blue equaled neurons clustered together on all runs whereas lighter shades proportionally reflected lower degrees of repeatable clustering. Robust clustering was evidenced by clear blue “blocks” of neurons along the matrix diagonal following ordering of cells by average linkage hierarchical clustering (HC) applied to this matrix. The stability of clustering was quantified by a cophenetic correlation coefficient computed between neural distances obtained from the consensus matrix and the dendrogram structure from HC.

### Analysis of neural population responses to taste and somatosensory stimuli

PB neurons recorded from different mice were combined into a hypothetical neural population to explore associations between stimulus responses across neurons. Similarity between responses was quantified pairwise using Spearman’s rank-order correlation coefficient. This non-parametric coefficient was used to accommodate skew and lack of normality in response data across cells (see Figure 3).

Correlations were visualized using colormap matrices and multidimensional scaling (MDS; Kruskal and Wish, 1978). For MDS, correlations between stimulus pairs were subtracted from 1 for conversion to proximities. MDS recovered these proximities as visualizable distances in a low-dimensional coordinate space. Stimuli that produced similar response patterns across neurons were plotted close together in this space. Dissimilar responses were positioned apart. To avoid local minima, MDS was repeated 50 times using random starting configurations. The MDS run that produced the lowest stress, which gauged the badness-of-fit of the reduction to the data, was used for final interpretation. Three-dimensional MDS was computed albeit only the first two dimensions were visualized for simplicity. MDS was performed using the *mdscale* function in MATLAB.

Neural population responses to the stimulus set were also explored using a self-organizing map (SOM). The SOM is an artificial neural network technique that can learn the structure of multivariate data and visually represent associations between data objects by their arrangement on a low-dimensional grid (Kohonen, 2013). We found that the SOM captured correlations between PB responses to the stimuli generally like MDS. However, the SOM additionally accounted for differences in stimulus response levels, which can elude correlation coefficients (Wilson et al., 2012; Schober et al., 2018).

In brief, the SOM algorithm projected the across-neuron response to each stimulus onto nodes of a 5 × 4 hexagonal grid; similar results were obtained with different grid sizes. Stimulus response data for each PB neuron were normalized between 0 and 1 prior to computing the SOM. Each grid node contained a weight (prototype) vector of the same length as the number of PB cells, and the algorithm found for each stimulus the node with weights closest to the across-neuron response. Using unsupervised learning, node weights were iteratively updated to bring the best-matching node for each stimulus, and neighboring nodes, closer to the stimulus response vector. The result of this process is that stimuli that cause similar PB responses best activate nearby nodes on the SOM grid. The visual topology of the best-matching nodes summarized the structure of stimulus activity across PB neurons. The SOM was applied using the SOM Toolbox (http://www.cis.hut.fi/somtoolbox/) version 2.1 in MATLAB. Batch training and a Gaussian neighborhood function were used.

Finally, principal component (PC) analysis was applied to visualize the distribution of PB neurons ordered by sensory tuning. Prior to this analysis, stimulus responses by each cell were standardized through division by the standard deviation of all responses and subtraction from the cell’s mean response. PC analysis was performed using the *pca* function in MATLAB.

### Behavioral measures

Methods to analyze brief-access licking data generally followed our prior work (Lemon et al., 2019). Briefly, median licks to each tastant concentration were computed across all test days, ignoring non-sampled trials. For individual mice, their licking responses to taste stimuli were converted to lick ratios, calculated as median stimulus licks divided by median licks to room temperature water. Trials for water (0 mM) were randomly interleaved into each taste stimulus presentation block during daily tests.

Lick ratios accommodate potential individual differences in responding and standardize data for comparisons. A lick ratio of 1 indicates equivalent licking to the stimulus and room temperature water (indifference). A lick ratio greater than 1 indicates more licks to (i.e., a preference for) the stimulus. Lick ratios less than 1 reflect reduced stimulus licks and are proportional to aversion.

### General analyses

Confidence intervals for medians and means were bootstrapped using the MATLAB function *bootci* with 1000 resamples. For some analyses, we used a Poisson statistical method to determine if stimulus response rates by individual cells were significantly larger than baseline activity, as described (Wilson and Lemon, 2014). Tests of the null hypothesis that equal frequencies of neurons responded across stimulus and experimental conditions were carried out using χ^2^ goodness-of-fit tests. χ^2^ also evaluated independence/association between variables. Between- and within-group comparisons involved parametric methods including *t*-tests or ANOVA. Non-parametric methods were used in several cases where data showed skew. These analyses were caried out using SPSS (version 27, IBM) or MATLAB. Normality of data samples was addressed using the Jarque-Bera goodness-of-fit test in MATLAB. Our criterion for statistical significance (α) was 0.05. When multiple/post-hoc tests were performed, *p*-levels were evaluated for significance using a false discovery rate (FDR) control procedure (Benjamini and Hochberg, 1995).

All data plots and graphs were generated using standard routines and custom code in MATLAB. Raw electrophysiological sweeps and thermal traces were exported from Spike2. Final figure configurations were produced using Illustrator (version 22.1, Adobe, San Jose, CA).

## Results

We generated TRPV1-ChR2 mice, which express ChR2 and EYFP in TRPV1-lineage fibers, by crossing TRPV1^Cre^ mice with the Ai32 Cre-reporter line. Ai32 mice express a ChR2/EYFP fusion protein following exposure to Cre recombinase. We made neurophysiological recordings from PB neurons in TRPV1-ChR2 mice and used an optrode to excite the terminals of TRPV1-lineage afferents that communicated with Vc neurons projecting to PB cells (Figure 1A). We observed that the spinal trigeminal tract near the orosensory dorsal Vc in TRPV1-ChR2 mice was heavily populated by ChR2-positive afferents (Figure 1B), consistent with prior observations of TRPV1-lineage fibers coursing this tract (Cavanaugh et al., 2011a; Mishra et al., 2011).

### ChR2 expression in the trigeminal tract was regulated by Cre in TRPV1-lineage fibers

In initial experiments, we used *in vivo* optrode recordings to show that pulses of blue laser light delivered to superficial tissues above the dorsal Vc could reliably excite Vc cellular activity in TRPV1-ChR2 mice (Figure 2D). Control experiments in Cre-negative Ai32 mice verified that in the absence of Cre recombinase, laser pulses had no effect on Vc cellular activity (Figure 2H). Fluorescence antibody amplification confirmed there was no expression of ChR2/EYFP in the trigeminal tract of Cre-negative Ai32 mice (Figure 2E-G). Notably, the same immunohistochemical procedures applied to detect Cre-directed fluorescence in tissues derived from TRPV1^Cre^ mice, where Cre recombinase is present in TRPV1-lineage fibers, revealed dense and focused labeling of neural processes within the spinal trigeminal tract, at rostrocaudal levels of the Vc (Figures 2A-C).

**Figure 2.**
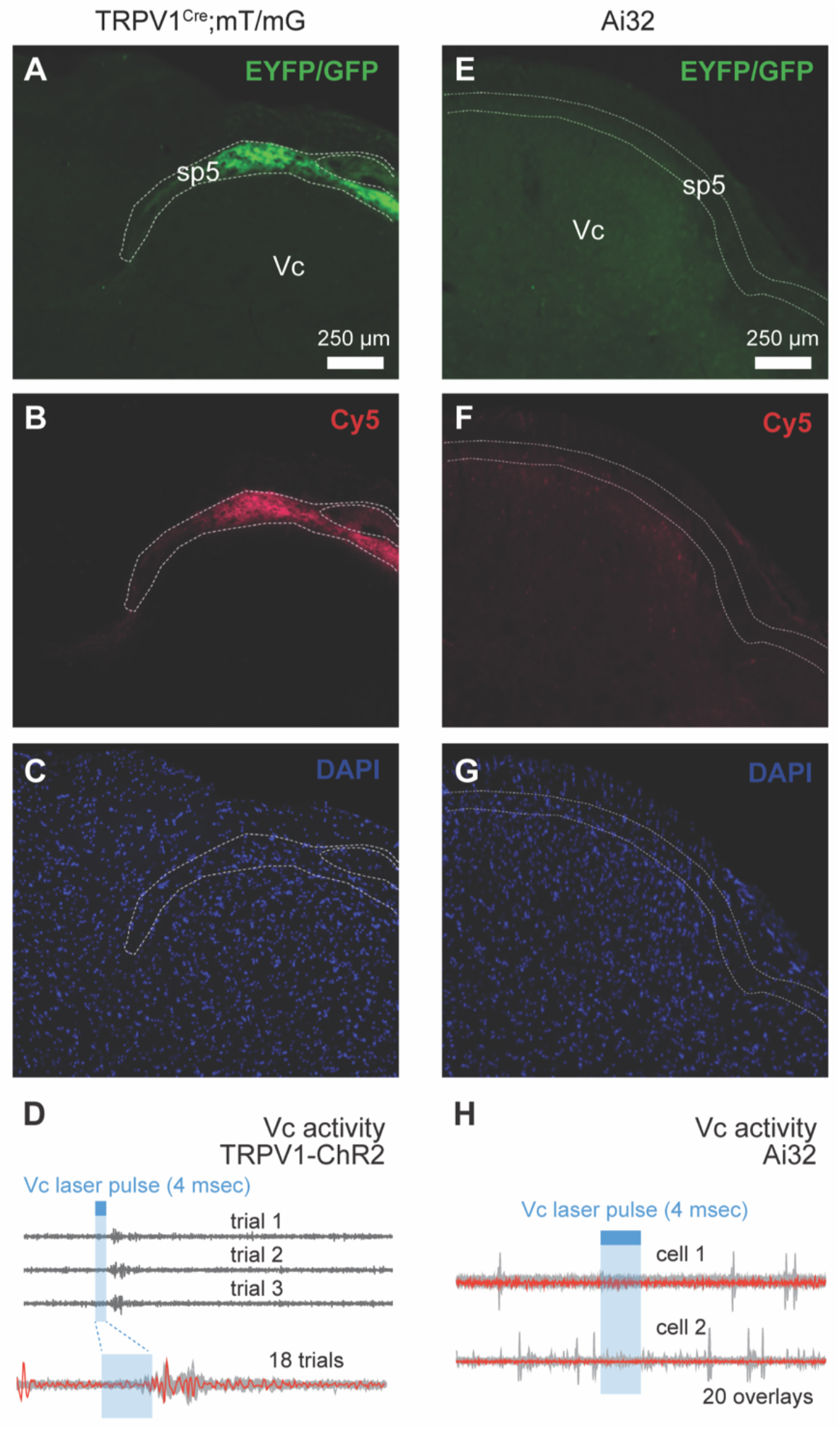
ChR2 expression was not leaky in Ai32 mice. A) Coronal brain section showing GFP labeling of TRPV1-lineage processes in the spinal trigeminal tract (sp5) of TRPV1^Cre^;mT/mG mice. These mice served as a positive control. B) Cy5 labeled antibody staining against EYFP/GFP in panel A. C) DAPI labeling of cell nuclei in the section in A and B. D) Example electrophysiological recordings of Vc activity in one TRPV1-ChR2 mouse. Laser pulses applied to the medullary surface above the orosensory Vc reliably stimulated multiunit activity. Red trace shows an example sweep for the overlay of responses across 18 trials. E) Coronal section showing no EYFP labeling of fibers in the sp5 of Cre-negative Ai32 mice, which express a ChR2/EYFP fusion protein only following exposure to Cre. F) Cy5 labeled antibody staining against EYFP/GFP applied the section in E yields no labeling, confirming ChR2/EYFP was not expressed. G) DAPI labeling showing cell nuclei in the section in E and F. H) Electrophysiological recordings of Vc unit activity in an Ai32 mouse. Laser pulses applied to the medullary surface above the orosensory Vc did not excite Vc neurons in this mouse line.

These results confirm that laser effects found in TRPV1-ChR2 mice were directed by Cre-induced expression of ChR2 in TRPV1-lineage afferents. Additionally, we made recordings of Vc activity in mice that expressed EYFP, but not ChR2, in TRPV1-lineage fibers but did not observe photoexcitatory responses (data not shown), implying laser effects were indeed ChR2 dependent.

### Trigeminal TRPV1-lineage fibers communicate with parabrachial taste neurons

PB neurons were sampled based on sensitivity to taste stimuli, with many of these cells also responding to changes in oral temperature (Figure 3). Relatedly, we encountered PB gustatory neurons that spiked to electrical and laser pulses directed to the orosensory dorsal Vc during neurophysiological recordings in TRPV1-ChR2 mice (Figure 1C-E). This implied PB taste neurons received input from upstream Vc circuits that included Vc neurons supplied by TRPV1-lineage afferents. We evaluated the frequency of this effect using electrophysiological data from 94 PB neurons that responded to temperature-controlled taste stimuli presented to the whole mouth (Figure 4A inset). Electrical pulse stimulation of the orosensory Vc significantly excited the majority (*n* = 62, 66%) of these cells (test of the null hypothesis that the activated proportion was chance, χ^2^ = 9.6, *df* = 1, *p* = 0.002; Figure 4A), agreeing with our previous study (Li and Lemon, 2019). Here, we found that optogenetic-assisted photoexcitation of TRPV1-lineage fibers arriving at the Vc caused responses in 87% (*n* = 54) of PB neurons that responded to Vc shocks (Figure 4A).

**Figure 3.**
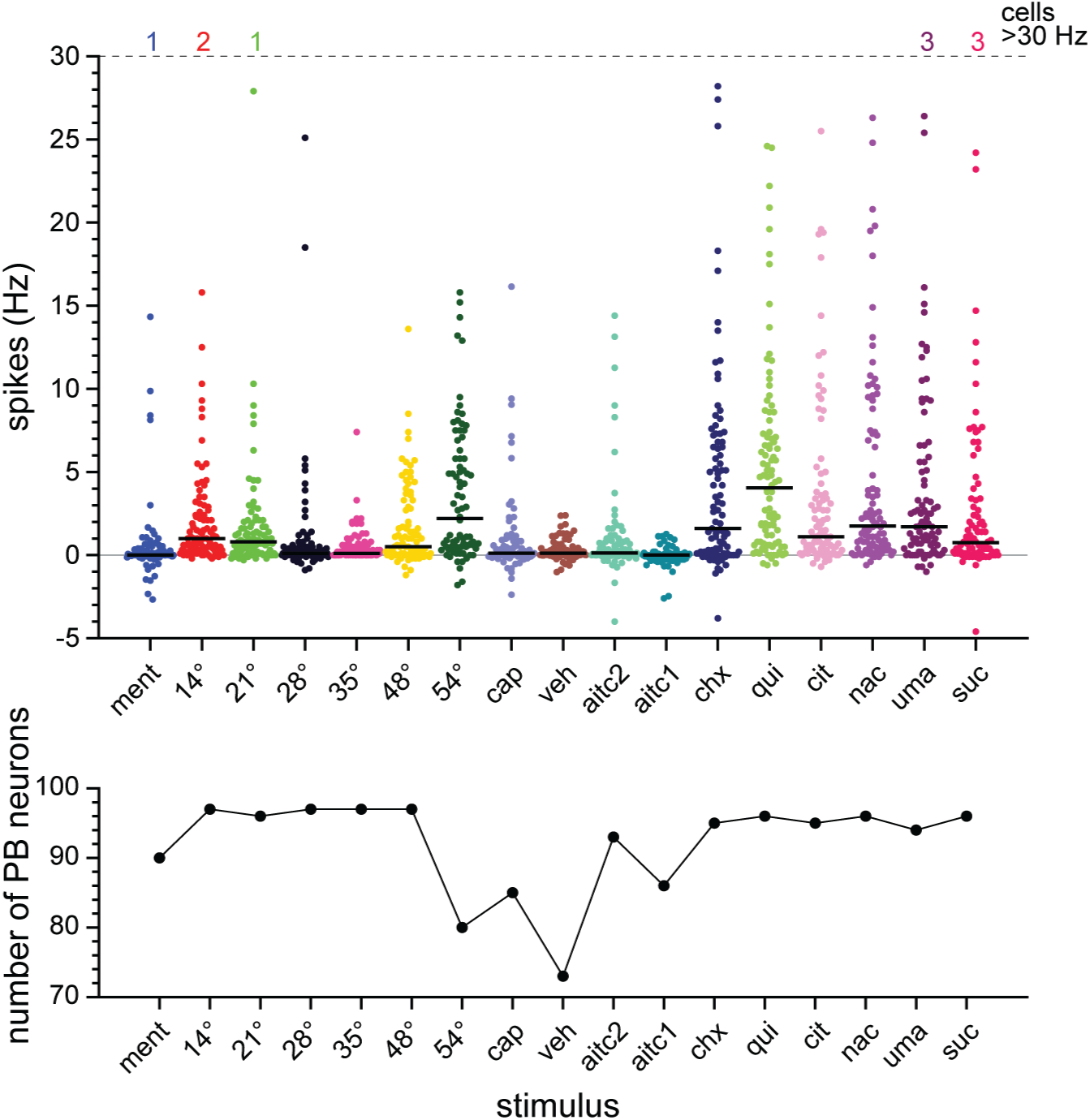
Stimulus responses by all sampled PB neurons. Top panel shows distributions of responses (circles; in spikes per sec) to each stimulus by individual cells. Horizontal bars give distribution medians. Integers at the top of the plot denote the number of cells that showed responses >30 Hz for select stimuli. On the x-axis, temperatures (in °C) represent oral stimulus temperatures on thermal ramps/trials (Figure 1F). Other abbreviations: ment, 1.28 mM (-)-menthol (delivery temperature: 28°C); cap, 1 mM capsaicin (room temperature); veh, capsaicin vehicle-only solution (room temperature); aitc2, 1 mM allyl isothiocyanate (35°C); aitd, 0.1 mM allyl isothiocyanate (35°C); chx, 0.1 mM cycloheximide (28°C); qui, 10 mM quinine-HCI (28°C); cit, 10 mM citric acid (28°C); nac, 100 mM NaCl (28°C); uma, umami mixture: 100 mM mono-K+ glutamate & 10 mM inosine 5’-monophosphate (28°C); sue, 500 mM sucrose (28°C). Bottom panel plots the number of neurons in each distribution. Cell numbers vary across stimuli as neurons did not always remain isolated for all stimulus tests. The number of neurons for capsaicin vehicle-only trials includes cells that also completed the blank control trial (see methods), which was used to correct neural activity on vehicle-only trials.

**Figure 4.**
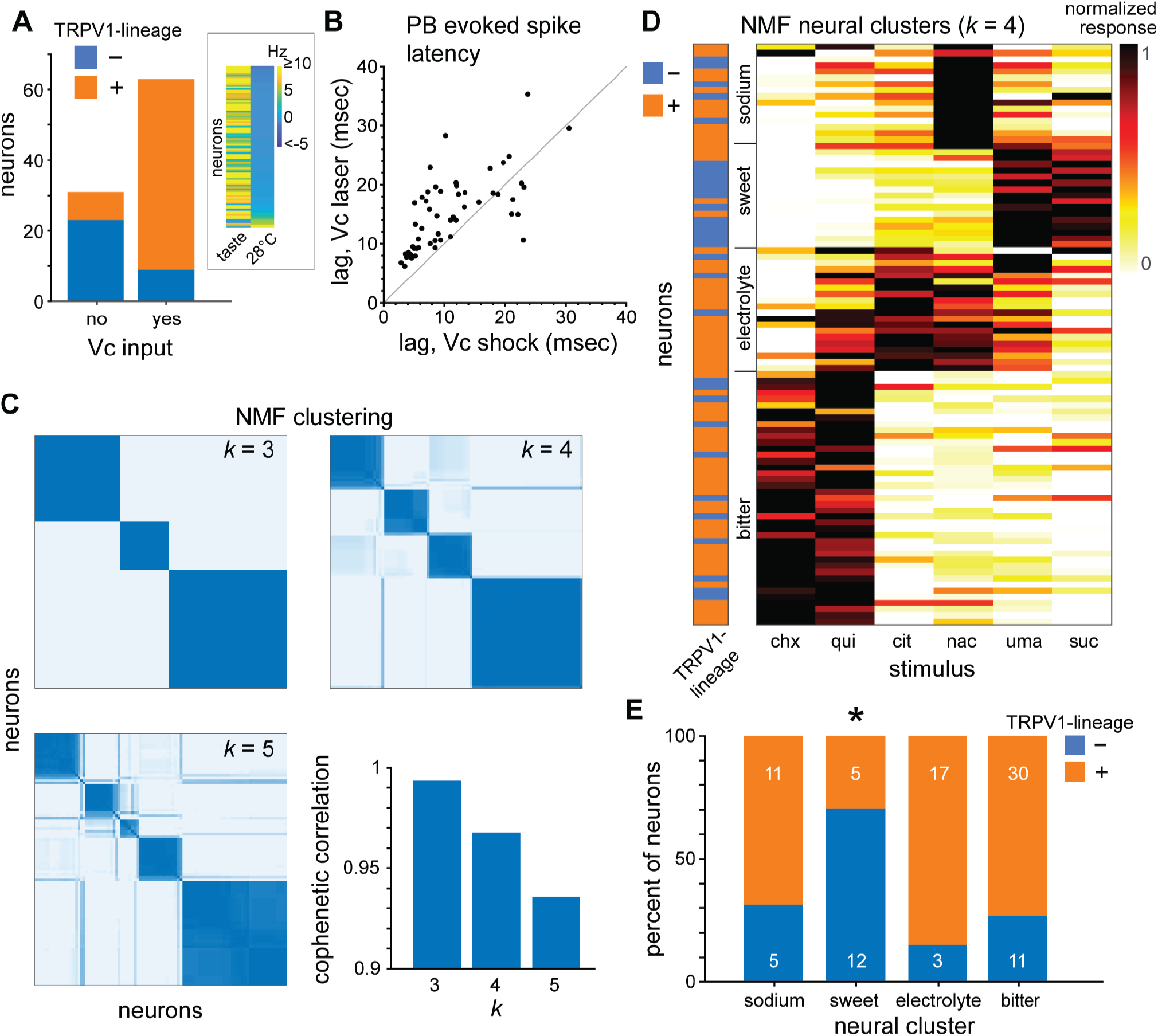
Contact from TRPV1-lineage afferents associates with gustatory tuning. A) The majority of PB neurons that responded to electrical stimulation of the Vc (input: yes) were TRPV1-lineage positive. Inset heatmap shows the highest taste response and the vehicle (28°C) response for all PB cells. B) In TRPV1-lineage positive neurons (circles), response latencies were longer to Vc laser pulses (y-axis) than shocks (x-axis; t-test, *p* = 0.000001). C, NMF identified 4 clusters of PB neurons (n = 94) from gustatory tuning. Consensus matrices show reliability of clustering for 3 to 5 groups (*k*). Deep blue cells mark neurons (axes) with reproducible clustering over many NMF runs. Lighter shades proportionally reflect neurons with inconsistent clustering, which frequently appear along the diagonal at *k* = 5. Thus, *k* = 4 supported stable clustering of the most groups. Cophenetic correlation quantifies the reliability of clustering. D, Heatmap shows normalized responses to taste stimuli and TRPV1-lineage input status for PB neurons in the four NMF clusters. For each neuron, taste responses are normalized across stimuli to make the smallest response 0 and the largest response 1, as represented by the color map legend. Taste stimuli are abbreviated as in Figure 3. E, Compared to other clusters, fewer PB sweet neurons were TRPV1-lineage positive (*, *x*^*2*^ test, *P =* 0.04). Integers on the percentage bar for each cluster give the actual numbers of TRPV1-lineage positive (upper orange section) and negative (lower blue section) neurons.

A significant association emerged between PB neuronal sensitivity to electrical (non-cell-type selective) and light (TRPV1-lineage specific) stimuli directed to oral Vc circuits (test of the null hypothesis of no association, χ^2^ = 33.2, *df* = 1, *p* < 0.001). Specifically, PB neurons that responded to Vc electrical shocks were more likely to fire to Vc laser pulses and meet criterion for classification as TRPV1-lineage positive cells (Figure 4A). Cellular latencies to respond to Vc electrical and light stimuli were positively correlated (Spearman’s rank-order correlation coefficient = +0.72, *p* < 0.001), with longer latencies found on laser pulse trials (paired *t*-test on normally distributed latency differences, *t*_52_= 5.5, *p* = 0.000001; Figure 4B).

Overall, these data show that neural information from a genetic type of somatosensory and nociceptive fiber is relayed by the Vc to PB neurons that respond to taste sensations. TRPV1-lineage positive gustatory neurons frequented the lateral PB nucleus including its external subregion (Figure 1H). Other studies have described gustatory activity in lateral PB areas including the external capsule (Yamamoto et al., 1994; Geran and Travers, 2009; Tokita et al., 2014; Tokita and Boughter, 2016; Jarvie et al., 2021), with the present and our prior work (Li and Lemon, 2019) identifying trigeminal projections reach taste cells in this PB region.

### Receipt of TRPV1-lineage input by parabrachial taste neurons is associated with gustatory tuning

A majority, but not all, of the 94 PB neurons responded to photoexcitation of TRPV1-lineage fibers. Thus, we used non-negative matrix factorization (NMF) to cluster cells by their responses to taste stimuli to study if gustatory tuning was related receipt of TRPV1-lineage input. NMF found four major clusters of PB neurons (Figure 4C). The taste profiles of neurons within these clusters were like groups reported in other studies of gustatory processing in the mouse PB area (Tokita and Boughter, 2012; Tokita et al., 2012; Tokita and Boughter, 2016; Li and Lemon, 2019). NMF clusters included neurons tuned to 1) sodium, 2) appetitive sucrose and the umami mixture (referred to as sweet cells for convenience), 3) electrolyte taste stimuli including citric acid, or 4) the bitter/toxic taste chemicals quinine and cycloheximide (Figure 4D). Notably, neural clusters were based on taste stimulus identity and not response intensity. Firing rates to strongly effective/best stimuli did not differ across NMF clusters (Independent samples Kruskal Wallis test, χ ^2^ = 5.2, *df* = 3, *p* = 0.2; Figure 5) whereas gustatory tuning did (Figure 4D).

**Figure 5.**
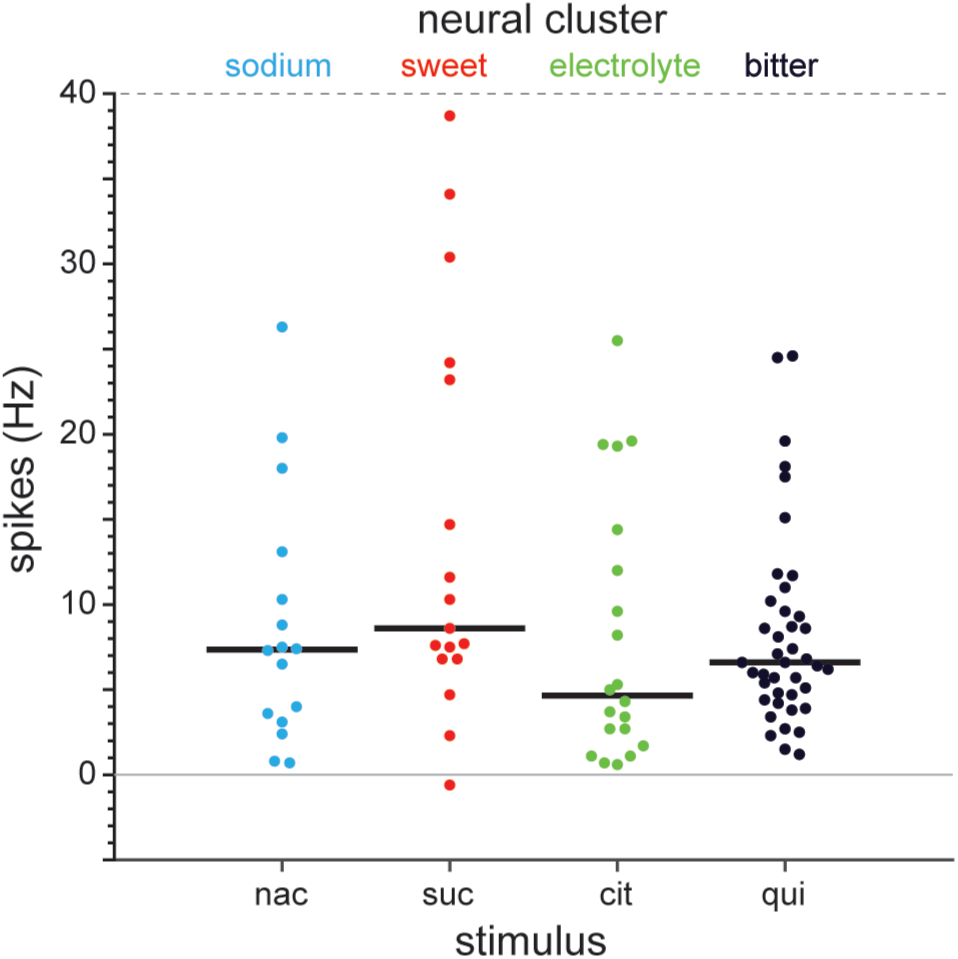
Neurons in different gustatory clusters show equivalent firing to their strongly effective/best stimuli. Plot shows distributions of responses (circles; in spikes per sec) to strongly effective/best taste stimuli (bottom x-axis) by individual cells in each NMF cluster (top x-axis). Horizontal bars give distribution medians. Responses rates to NaCl (nac) by sodium cells, sucrose (sue) by sweet neurons, citric acid (cit) for electrolyte neurons, and quinine (qui) in bitter cells did not differ (Kruskal Wallis test, *p =* 0.2). This result implied neural clusters were based on stimulus identity and not response intensity. Concentrations/temperatures of taste stimuli are as in Figure 3.

Photoexcitation of TRPV1-lineage fibers arriving at the orosensory Vc caused significant responses in the majority of sodium (11 of 16, 69%), electrolyte (17 of 20, 85%), and bitter (30 of 41, 73%) PB neurons (Figure 4D, 4E). However, only a minority of sweet neurons (5 of 17, 29%) oriented to sucrose and the umami mixture responded to photoexcitation of TRPV1-lineage fibers (Figure 4D, 4E). Frequency analysis applied to sodium, sweet, and electrolyte neurons, which had similar sample sizes (Figure 4D) to mitigate bias, revealed that the reduced number of TRPV1-lineage positive cells within the sweet PB cluster was significant (test of the null hypothesis that across clusters, equal numbers of neurons were TRPV1-lineage positive, χ ^2^ = 6.5, *df* = 2, *p* = 0.04; Figure 4E).

These results suggest that gustatory tuning partly predicts whether PB taste neurons receive input from trigeminal TRPV1-lineage fibers. The observed pattern of convergence also appears to inversely relate to mouse behavioral preferences for tastes. TRPV1-lineage afferents only infrequently contacted PB gustatory neurons oriented to sucrose and umami stimuli, which mice show strong preferences for in brief-access fluid licking tests (Ellingson et al., 2009; Saites et al., 2015). Accordingly, we found that thirst-motivated C57BL/6J mice (*n* = 6) preferred to lick the 500 mM concentration of sucrose tested on PB neurons, and also 1000 mM sucrose, over water in a brief-access setting (FDR-corrected *t*-tests on normal data comparing mean lick ratios to 1, *t*_5_ > 3.5, *p* < 0.02; Figure 6A).

**Figure 6.**
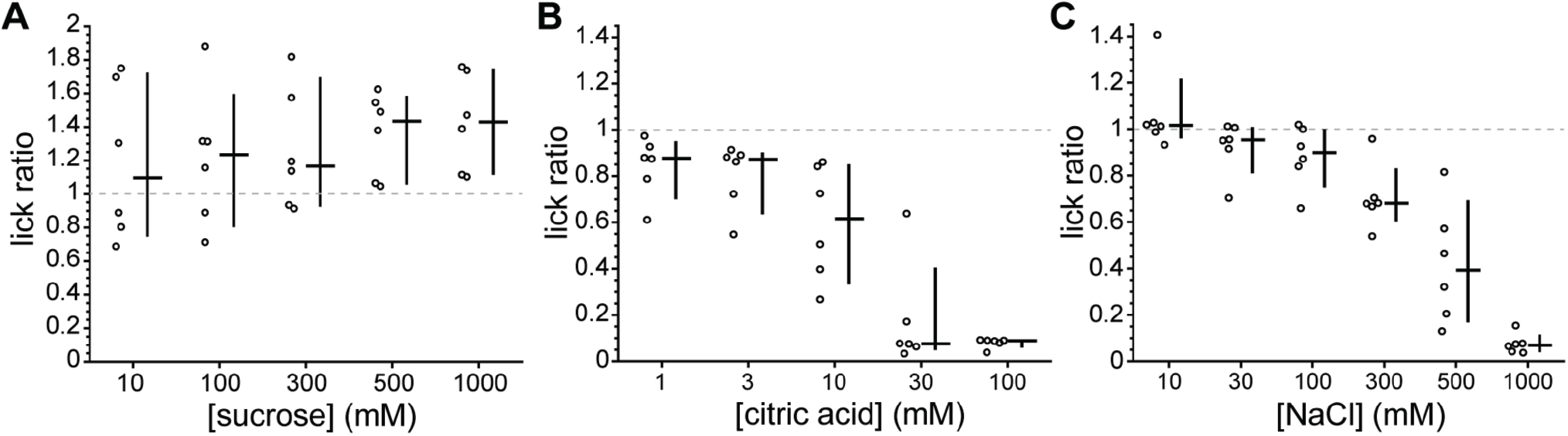
Mouse oral behavioral preferences for select taste stimuli. Each panel plots distributions of lick ratios (licks to stimulus - licks to water) measured by brief-access fluid lickometry for individual concentrations of a room temperature taste stimulus. All data in this figure were collected from one group of C57BL/6J mice (n = 6). For each concentration, circles represent median lick ratios for individual mice across all sampled trials/test days. Lines to the right of each lick ratio distribution give its median (horizontal bar) and the bootstrapped 95% confidence interval of the median (vertical bar). A) Lick ratios for a 10 to 1000 mM concentration series of sucrose. Mice showed more licks to 500 and 1000 mM sucrose than water (lick ratios > 1, t-tests, *p* < 0.02). B) Lick ratios for a 1 to 100 mM concentration series of citric acid. C) Lick ratio data for a 10 to 1000 mM concentration series of NaCl. For citric acid and NaCl, lick ratios shifted from indifference compared to water (lick ratios near 1) to aversion (lick ratios near 0) with rising concentration (Friedman’s ANOVAs, *p* < 0.0004). In general, mice showed only a moderate reduction in licking and indifference, respectively, to 10 mM citric acid and 100 mM NaCl tested on PB neurons.

Preference for sucrose and umami is unique compared to the other presently tested taste stimuli, which can cause behavioral aversion at the present or higher concentrations and best stimulated PB neurons that mostly received TRPV1 afferent input. Prior data establish that mice avoid milli- and micromolar concentrations of the bitter stimuli quinine and cycloheximide in brief-access tests (Long et al., 2010; Lemon et al., 2019). Although there are few comparable data for citric acid, we found that the 10 mM concentration tested on PB neurons caused only a moderate reduction in brief-access licking in mice compared to water. However, higher concentrations of citric acid substantially decreased licking (Friedman’s ANOVA by ranks, χ ^2^ = 21.07, *df* = 4, *p* = 0.0003) and induced clear orosensory aversion to near-zero licks (Figure 6B). This trend parallels mouse licking behaviors toward NaCl, which largely evoked indifference in licking compared to water at lower millimolar concentrations, including 100 mM tested on PB neurons, but reduced licks with rising concentration (Friedman’s ANOVA by ranks, χ^2^ = 25.5, *df* = 5, *p* = 0.0001) and aversion at molar levels (Figure 6C; Glendinning et al., 2002).

Overall, TRPV1-lineage somatosensory fibers appear to only sparingly communicate with PB sweet cells that respond best to innately preferred tastes. Importantly, the grouping of neurons by taste was computed independently of testing for TRPV1-lineage input status, with significantly fewer sweet cells responding to optical activation of TRPV1 fibers.

### TRPV1-lineage input is associated with oral thermal activity in parabrachial taste neurons

TRPV1-lineage fibers comprise primary thermosensory neurons activated by cold or heat (Mishra et al., 2011). Oral temperatures can excite PB taste neurons that respond to electrical stimulation of the Vc (Li and Lemon, 2019). Thus, we investigated if TRPV1-lineage positive PB neurons were temperature sensitive. For thermal stimuli, the oral cavity was bathed with purified water that was rapidly stepped from physiological temperature (35°C) to cool (28°, 21°, and 14°C), neutral (35°C), and noxious hot (48° and 54°C) temperatures inside the mouth on discrete trials (Figure 1F).

Inspection of spike trains revealed that many TRPV1-lineage positive PB neurons responded to the cold and heat limits of 14° and 54°C (Figures 1F, 7A). These temperature limits stimulated significant firing in the majority of TRPV1-lineage positive cells (14°C: 35 of 66 neurons, 53%; 54°C: 37 of 53 neurons, 70%; Figure 7B). Increasing numbers of TRPV1-lineage positive PB neurons were recruited as temperature steps approached and reached either 14° or 54°C (test of the null hypothesis that equal frequencies of neurons activated across temperatures excluding 35°C, χ^2^ = 16.7, *df* = 4, *p* = 0.002; Figure 7B). In contrast, only a reduced minority of TRPV1-lineage negative PB neurons responded to 14° and 54°C (Figure 7B), with the proportion of activated neurons invariant across the temperature steps (χ^2^ = 5.04, *df* = 4, *p* = 0.3).

**Figure 7.**
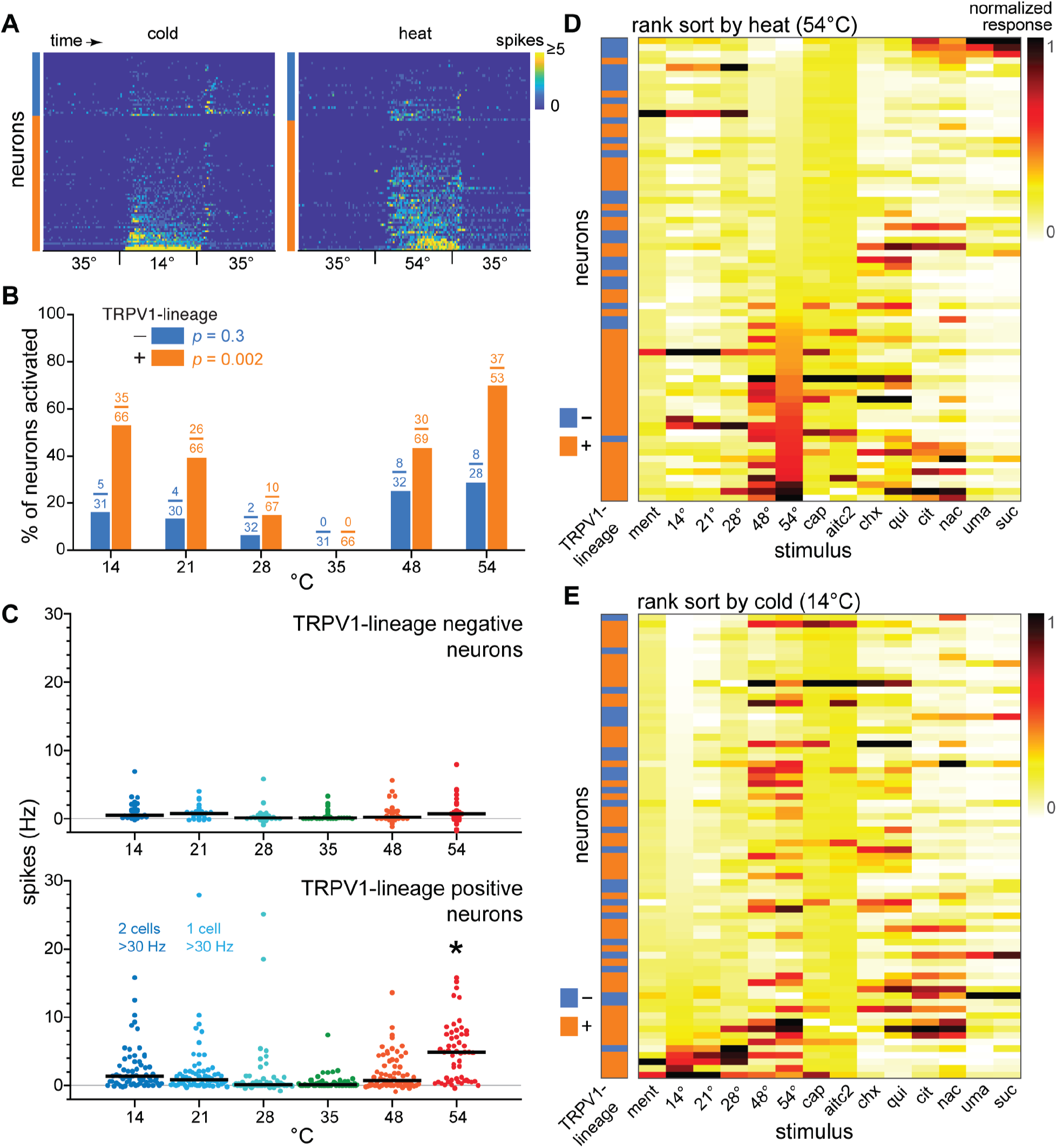
Temperature responses predominantly emerge in PB taste neurons excited by TRPV1-lineage afferents. A) Heatmaps show stacked peristimulus time histograms (spikes per 100 msec, legend) for PB neurons across all trials where 14°C (n = 97) or 54°C (n = 81) water was orally delivered for 5 sec. Color bar to the left of the y-axis of each plot marks the TRPV1-lineage positive/negative status of the cells (colors follow legend in B). B) Cooling or heating steps from 35°C excited increasing numbers of TRPV1-lineage positive (χ^2^, *p =* 0.002), but not negative (χ^2^, *p =* 0.3), PB neurons. Fraction above each bar gives the number of cells that showed significant excitation (numerator) over the total number of neurons analyzed. C) Responses (circles; in spikes per sec) to oral temperatures in PB neurons. Horizontal bars are medians. Responses to 54°C were larger in TRPV1-lineage positive than negative PB cells (*, Wilcoxon test, *p =* 0.0002). D) Heatmap shows normalized responses and TRPV1-lineage input status for PB neurons (*n* = 70) sorted in ascending order by their response to 54°C. For each stimulus, responses are normalized across neurons to make the smallest cell response 0 and the largest 1, as represented by the color map legend. See Figure 3 for stimulus abbreviations. E) Same as D, but neurons are sorted by their response to 14°C.

Analysis of firing rates found that responses to extreme heat at 54°C were larger in TRPV1-lineage positive than negative PB neurons (FDR-corrected Wilcoxon test, *p* = 0.00004; Figure 7C).

TRPV1-lineage positive neurons also appeared to show larger responses to noxious heating to 48°C and to the cold limit of 14°C. However, these differences in temperature activity against TRPV1-lineage negative cells did not survive FDR control of *p*-levels for multiple comparisons (Wilcoxon tests, *p* > 0.025). Notably, TRPV1-lineage positive neurons showed greater firing to 54° than 14°C (paired *t*-test with normally distributed response differences, *t*_52_ = 3.9, *p* = 0.0003; Figure 7C), which agrees with the lower spike density many of these cells displayed during the cold stimulus period (Figure 7A). The reduced responses to 14° and 54°C in TRPV1-lineage negative neurons did not differ (paired *t*-test with normally distributed response differences, *t*_26_ = 0.2, *p* = 0.8).

Finally, rank sorting PB neurons by their responses to 54° and 14°C revealed the largest responses to oral heating or cooling emerged nearly exclusively in TRPV1-lineage positive cells (Figure 7D, 7E). PB neurons that showed the highest activity to 54° and 14°C included cells that responded to tongue presentation of capsaicin (Figure 1G, 7D), which stimulates the heat nocisensor TRPV1, or whole mouth delivery of menthol (Figure 7E), which engages the cold receptor TRPM8. In adult mice, TRPV1 and TRPM8 are expressed by TRPV1-lineage afferents (Mishra et al., 2011).

The above results demonstrate that thermal activity in PB gustatory neurons is associated with input from TRPV1-lineage fibers. These data also represent an initial account of how the sensory properties of taste neurons are related to the actions of genetically defined somatosensory neurons.

### Cold and heat differentially stimulate parabrachial taste neurons

We noted that while the oral presence of heat ≥48°C could induce strong activity in PB neurons co-sensitive to tastes (Figure 7D), the largest responses to cold at 14°C emerged in PB cells that showed comparably weak responses to gustatory stimuli (Figure 7E). To explore this further, we reapplied NMF to cluster neurons that completed all temperature, chemesthetic, and taste trials (*n* = 70) by their responses to gustatory and thermal stimuli (Figure 8A). This analysis captured the clusters of PB taste neurons observed in the prior matrix factorization (Figure 4D) and also two additional clusters of predominantly TRPV1-lineage positive cells sensitive to oral cooling or heat (Figure 8B).

**Figure 8.**
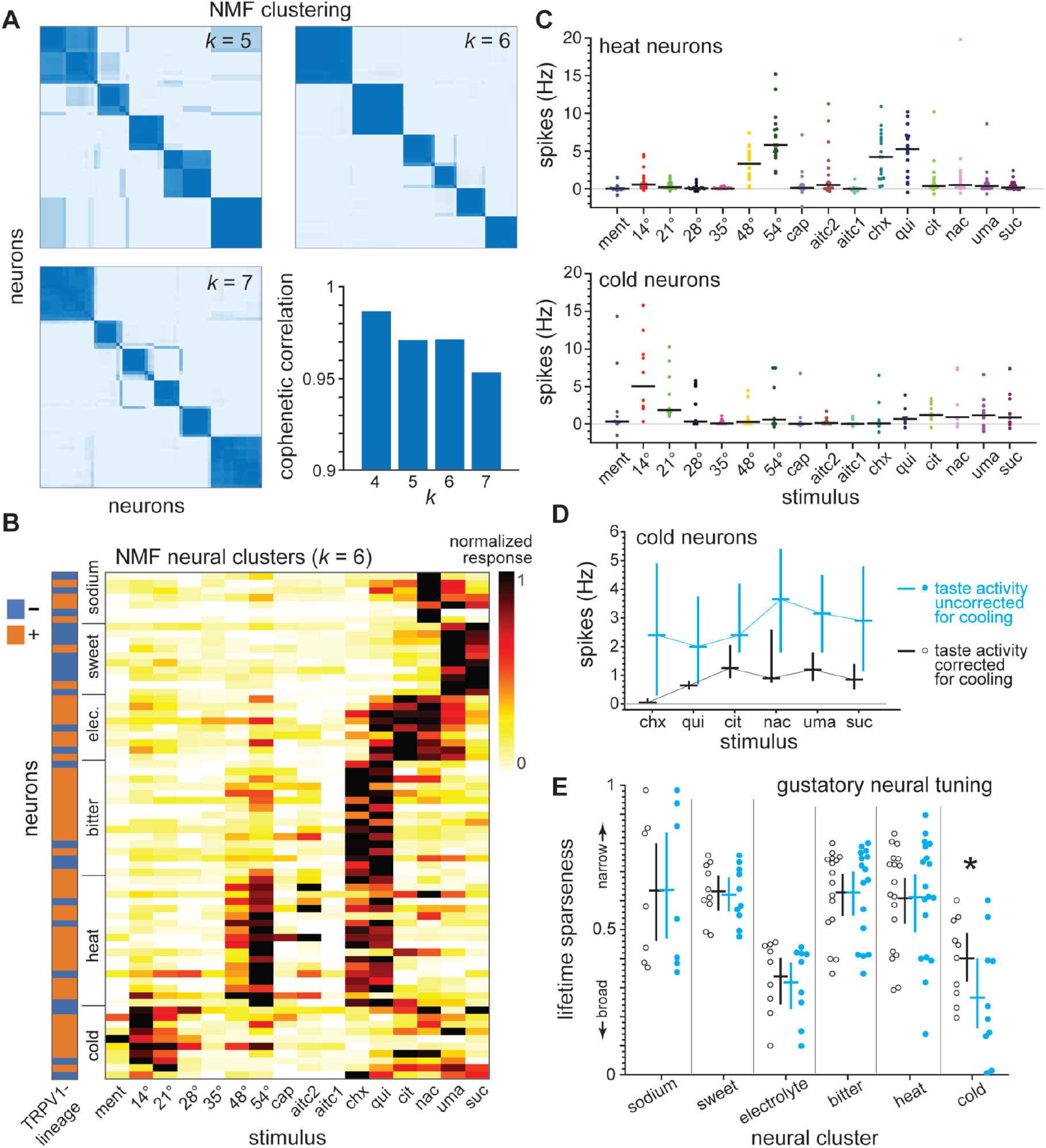
Heat and cold differentially excite PB gustatory neurons. A, NMF identified 6 clusters of PB neurons *(n =* 70) based on taste and temperature responses. Consensus matrices and cophenetic correlation coefficients show reliability of clustering for *k =* 5 to 7. Robust clustering (e.g., deep blue blocks of neurons along consensus matrix diagonal) was noted for *k =* 6, with performance decreasing with *k=7*. B, Heatmap shows normalized responses and TRPV1-lineage input status for PB neurons in the 6 clusters. For each cell, responses are normalized across stimuli to make the smallest response 0 and the largest 1, as per the legend. See Figure 3 for abbreviations. C, Distribution of responses (circles; in spikes per sec) to all stimuli by heat and cold neurons. Horizontal bars are medians. Heat neurons (*n* = 18) showed strong and equivalent (f-test, *p* = 0.1) responses to oral delivery of noxious heat at 54°C and bitter taste stimuli (cycloheximide [chx] and quinine [qui]). Cold neurons (*n* = 10) showed comparably low responses across taste stimuli and greater activity to oral cooling to 14°C (f-tests, *p* < 0.05). D, Median responses (horizontal bars) to taste stimuli across cold neurons calculated from cellular activity both corrected and uncorrected for spikes to the cooling ramp that accompanied taste delivery (legend). Vertical bars represent the bootstrapped 95% confidence interval of each median. The low, temperature-corrected responses to quinine, citric acid (cit), NaCl (nac), sucrose (sue), and the umami mixture (uma) were near equivalent. Across taste stimuli, responses broadly increased when activity was uncorrected for the cooling ramp of taste solutions. E, This increase in response breadth in cold neurons was unique and significant (*, two-way ANOVA post-hoc test, *p =* 0.00002).

Heat neurons showed strong responses to extreme heat at 54°C and the bitter taste stimuli quinine and cycloheximide (Figure 8B, 8C). In heat cells, bitter tastants induced responses of the same magnitude as 54°C (paired *t*-test comparing spikes to 54°C and the maximum response to cycloheximide or quinine, *t*_17_ = 1.6, *p* = 0.1; response differences were normally distributed).

In contrast, cold neurons could show larger responses to cool temperatures, such as 14°C, than to taste stimuli (FDR-controlled paired *t*-tests comparing activity to 14°C and each taste stimulus, *t*_9_ > 2.7, *p* < 0.05; response differences were normally distributed; Figure 8B, 8C). Cold neurons displayed median temperature-corrected responses to taste stimuli that were just above zero (Figure 8C). Although low, the generally flat responsiveness of cold neurons across gustatory stimuli (Figure 8D) rendered these cells some of the more “broadly tuned” across tastes by conventional metrics (Figure 8E). Among neural clusters, cold neurons also uniquely increased their breadth of responsiveness across taste stimuli when gustatory responses were uncorrected for the cooling ramp that accompanied taste delivery (FDR-controlled post-hoc test, *p* = 0.000004; neural cluster × thermal correction status interaction on lifetime sparseness, *F*_5,64_ = 4.3, *p* = 0.002; Figure 8E).

While the majority of TRPV1-lineage positive PB neurons significantly responded to oral cooling to 14°C or noxious heat at 54°C (Figure 7B), only noxious heat induced firing rates comparable to temperature-corrected taste activity in PB gustatory cells (Figure 8B, 8C). The strongest responses to cold temperatures emerged in TRPV1-lineage positive PB neurons with limited sensitivity to tastes. These “cold” neurons (Figure 8B-D) were possibly acquired due to responsiveness to the cooling (Figure 1E), but not chemosensory, component of taste stimuli. Nevertheless, because neural sampling was guided by sensitivity to taste solutions, our results imply that neural signals for oral heat and cooling differentially overlap with taste processing in PB circuits.

### Responses by parabrachial neurons capture hedonic relationships between taste and somatosensory stimuli

We observed certain trends in PB thermo-gustatory responses that appeared to convey hedonic features of taste and somatosensory stimuli. For instance, rank sorting neurons by temperature activity revealed the smallest responses to noxious heat at 54°C emerged in cells that showed the largest responses to the preferred tastes sucrose and umami (Figure 7D). On the other hand, noxious heat ≥48°C produced robust responses in PB neurons that strongly fired to aversive bitter tastes (Figure 8B, 8C).

To further explore stimulus relationships, we used multidimensional scaling (MDS) to visualize correlations between responses to all gustatory and somatosensory stimuli across the 70 PB neurons that completed all stimulus tests. MDS produced a coordinate space that used proximity to visually represent correlations between population responses to the stimuli, with positively correlated responses placed near one another. Along this line, PB responses to noxious heat ≥48°C and the bitter tastants quinine and cycloheximide were clustered together in MDS space (Figure 9A). MDS separated heat and bitter responses from oral cooling stimuli and moderate (100 mM NaCl and 10 mM citric acid) and preferred (sucrose and the umami mixture) tastes (Figure 9A). Notably, preferred tastes were located the furthest away from nociceptive and bitter stimuli along the first dimension of the MDS space, which reflected wide differences in responses to these stimuli across PB neurons.

**Figure 9.**
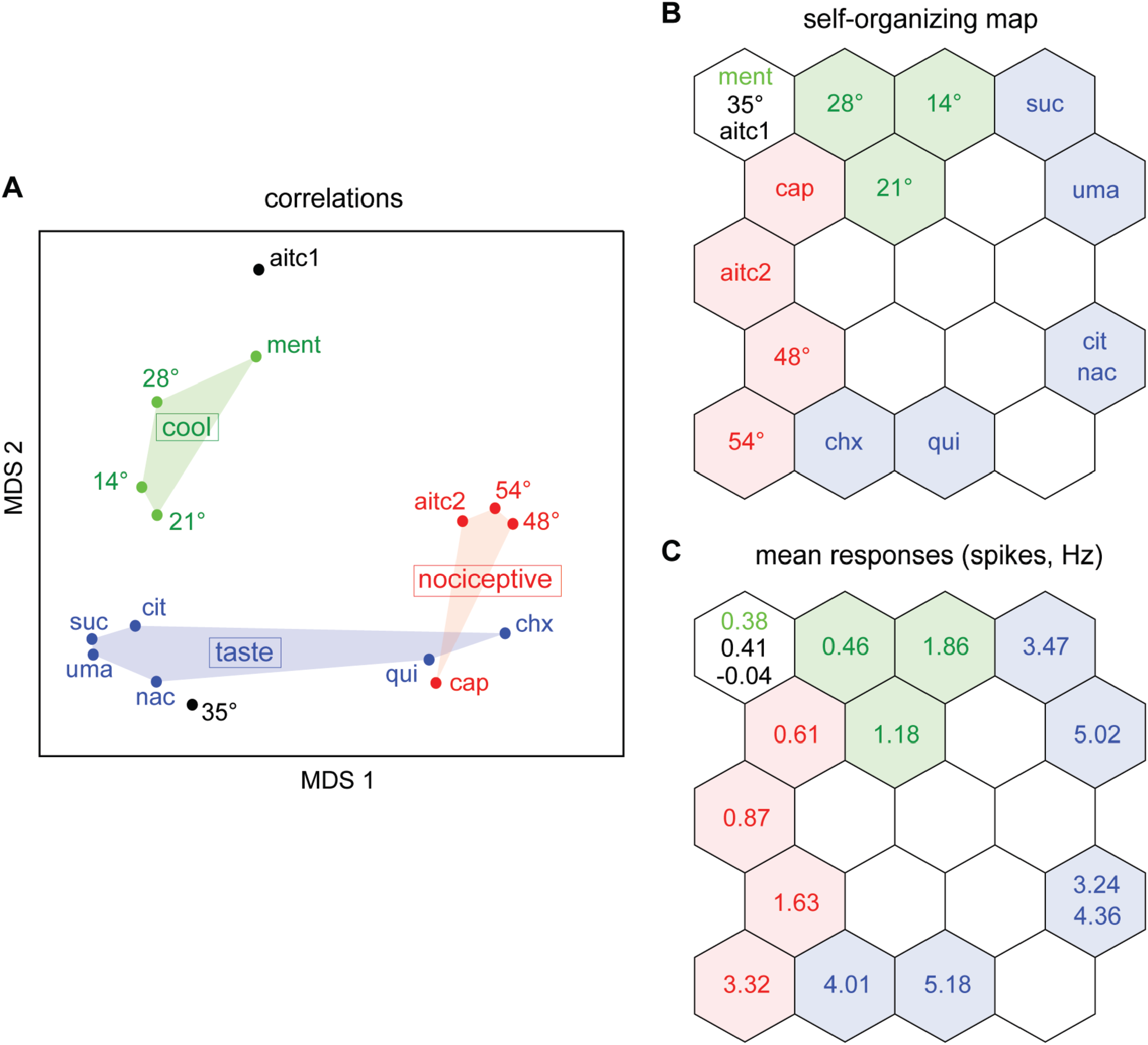
PB population responses distinguish aversive taste and nociceptive stimuli from other gustatory and somatosensory inputs. A) Multidimensional scaling (MDS) visualized the correlation structure of the responses to all stimuli across 70 PB neurons. Based on proximity in MDS space, bitter tastants (quinine [qui] and cycloheximide [chx]) induced PB responses that were positively associated with activity to noxious heat (48° and 54°C) and chemonociceptive stimuli (capsaicin [cap] and an elevated concentration of allyl isothiocyanate [aitc2]). See Figure 3 for all stimulus abbreviations. B) Similarity grid of stimulus responses across the 70 PB neurons produced by a self-organizing map (SOM). Following SOM training, stimuli that induce similar responses are associated with nearby nodes of the grid. Stimulus labels mark their best-matching node. Noxious heat (48° and 54°C) and bitter taste stimuli (chx and qui) best stimulated adjacent nodes in the lower-left corner of the grid, reflecting correlated activity. Notably, bitter and heat stimulated a region of the SOM grid opposing that engaged by preferred tastes like sucrose. C) In addition to recovering correlations, the SOM also ordered stimuli by response magnitude. Here, stimulus labels for best-matching nodes in panel B are exchanged for the mean response each stimulus evoked across PB neurons (e.g., chx = 4.01 Hz). Stimuli that induced weak responses (e.g., 35°C and a low concentration of allyl isothiocyanate [aitd]) engaged the same node in the upper-left corner of the SOM grid. With rising response levels, nociceptive agents (cap and aitc2) best stimulated nodes that tracked toward the SOM location for aversive heat and bitter stimuli. Cooling temperatures engaged SOM entries that, with rising response levels, tracked towards the best-matching unit for sucrose.

MDS positioned PB responses to the nociceptive agents capsaicin and an elevated concentration of AITC (1 mM) near bitter and heat stimuli in the coordinate space (Figure 9A). While some neurons did show strong firing to capsaicin and 1 mM AITC (Figure 1G), these responses were comparably sparse across PB cells (Figure 3). Nonetheless, the strongest responses to capsaicin and 1 mM AITC predominantly emerged in PB heat/bitter cells (Figure 8B, 8C), agreeing with the MDS solution. We found that a reduced concentration (0.1 mM) of AITC and 35°C water induced weak firing (Figure 3, 8B) and were scattered in MDS space (Figure 9A). This reflects the insensitivity of correlation coefficients to comparably low responses (Wilson et al., 2012; Schober et al., 2018).

These interpretations of the stimulus response structure were confirmed by a self-organizing map (SOM). The SOM recovered the similarity of PB responses to the bitter tastants and noxious heat, as reflected by their adjacent positioning on the SOM grid (Figure 9B). Responses to aversive heat and bitter stimuli excited a region of the SOM grid that opposed the location of preferred sucrose and umami stimuli, which followed the wide differences between these responses recovered by MDS. Moreover, the SOM recognized stimuli that produced weak responses across neurons, such as 35°C water and 0.1 mM AITC, and clustered them together (Figure 9B, 9C). With increasing response levels, 1 mM AITC and capsaicin tracked toward the heat/bitter cluster on the SOM grid (Figure 9B, 9C). Cooling temperatures tracked in a different direction with rising response levels, toward the grid location for preferred sucrose (Figure 9B, 9C).

Together, the results of MDS and the SOM suggest that the correspondence between PB responses to bitter tastants and noxious heat distinguishes their common aversive value from the hedonic features of other stimuli. We assessed correlations between responses to noxious heat, capsaicin, 1 mM AITC, and bitter tastants in TRPV1-lineage positive and negative PB neurons to explore how these cell subpopulations contribute to the clustering of aversive stimuli. In TRPV1-lineage negative neurons (*n* = 24), responses to the bitter tastants cycloheximide and quinine were uncorrelated (*p* > 0.01, evaluated under FDR control for multiple tests) with activity to nociceptive stimuli, including noxious heat ≥48°C and capsaicin (Figure 10A). In contrast, TRPV1-lineage positive PB neurons (*n* = 46) gave response to bitter tastants that did show various levels of significant positive correlation (*p* < 0.007, evaluated under FDR control) with nociceptive stimuli (Figure 10B). This trend was also observed across several random subsamples of 24 TRPV1-lineage positive neurons (e.g., Figure 10C). These subsamples equated the number of TRPV1 positive and negative cells to mitigate bias on significance levels imposed by different sample sizes. This exploratory analysis implies input from TRPV1-lineage fibers drives the association of nociceptive activity with aversive taste sensations conveyed by PB gustatory cells. The above also opens the possibility that TRPV1-lineage positive and negative PB neurons serve different functions for oral sensory coding.

**Figure 10.**
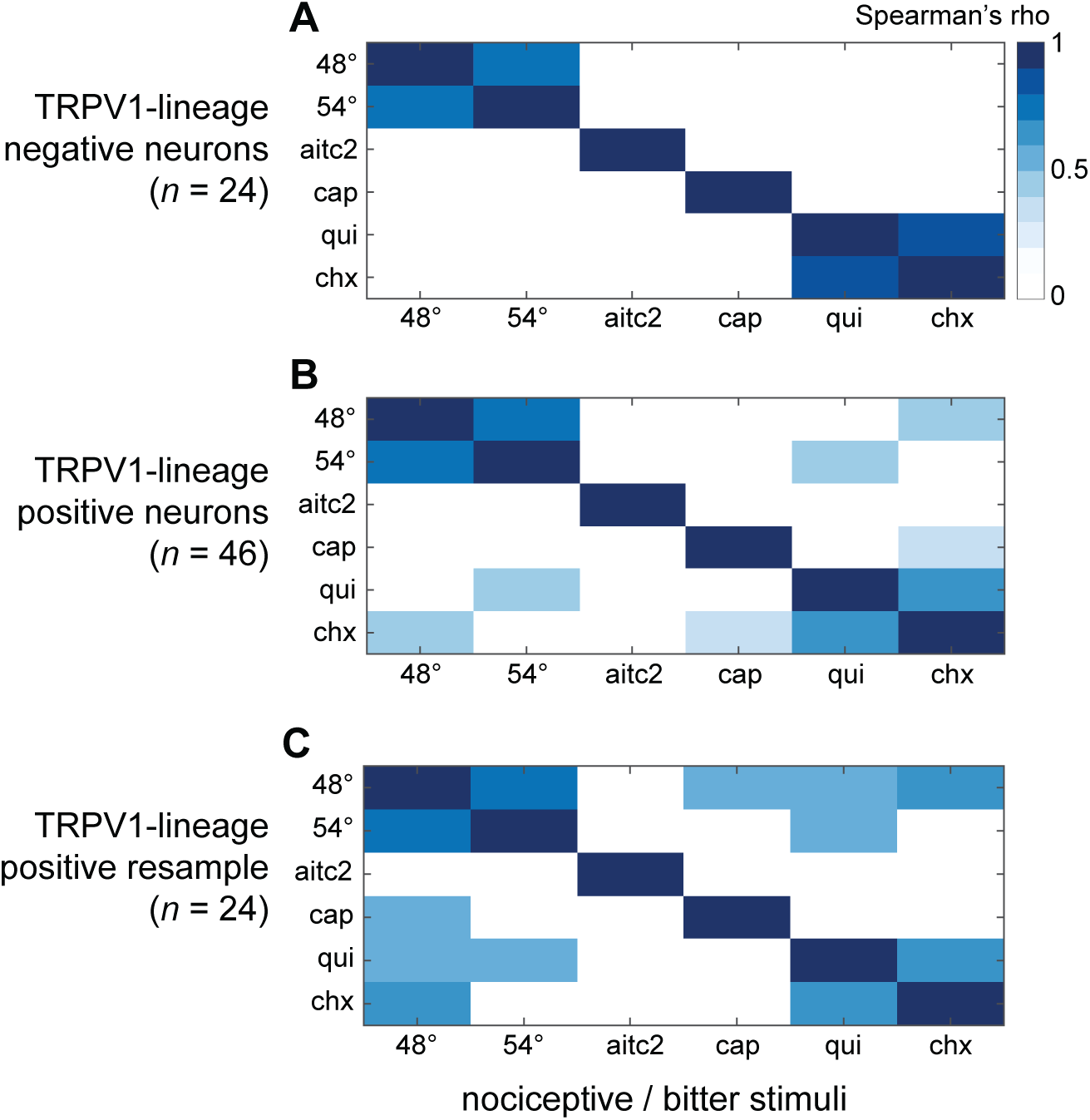
TRPV1-lineage positive and negative PB neurons capture different stimulus features. A) Correlation matrix for responses to bitter taste and nociceptive stimuli in TRPV1 -lineage negative PB neurons (n = 24). For all panels of this figure, white matrix entries represent non-significant correlations. Blue shades denote significant correlations (p < 0.02, evaluated using FDR control for multiple tests) and coefficients (color bar legend). ForTRPVI-lineage negative PB neurons, no significant positive correlations were found between their responses to bitter taste (quinine [qui] and cycloheximide [chx]) and nociceptive stimuli (48°C, 54°C, allyl isothiocyanate [aitc2], and capsaicin [cap]). B) Same as A, except for TRPV1-lineage positive cells *(n =* 46). C) Same as A and B, except the correlation matrix was computed for bitter taste and nociceptive responses by a random (without replacement) subsample of TRPV1-lineage positive PB neurons (*n* = 24) drawn to equate the numbers of TRPV1-lineage positive and negative cells. In panels B and C, various degrees of significant positive correlation emerged between responses to bitter taste and nociceptive stimuli by TRPV1-lineage positive PB neurons.

## Discussion

Using Cre-directed optogenetics and neurophysiology, we found that genetically defined thermosensory and pain fibers are components of trigeminal circuits that project to and excite mouse PB gustatory neurons. Further, this confluence of somatosensation with taste drove correlations between responses to aversive taste and nociceptive stimuli. The significance of these observations is discussed below.

### Thermal nociceptive fibers communicate with parabrachial taste neurons

We previously demonstrated that trigeminal neurons in the dorsal Vc project to taste neurons in the mouse PB nucleus and contribute to temperature responses in these cells (Li and Lemon, 2019). However, the cell types involved with this convergence were undefined. Here we show that TRPV1-lineage thermosensory and nociceptive fibers are primary neurons that drive trigeminal circuits projecting to PB taste neurons. Photoexcitation of TRPV1-lineage afferents arriving at the orosensory dorsal Vc frequently stimulated PB gustatory cells. PB neurons that responded to excitation of TRPV1 afferents displayed greater responsiveness to oral temperatures, including noxious heat, than TRPV1-lineage negative PB neurons. This implies TRPV1-lineage fibers contribute to thermal responses in PB orosensory and taste cells.

Our methods aimed to stimulate the terminals of TRPV1-lineage afferents that communicated with Vc neurons conveying orofacial sensations to PB circuits. Accordingly, PB neurons showed longer latencies to respond to optical excitation of TRPV1-lineage fibers than non-selective electrical stimulation of the Vc. While this agrees with photoexcitation of TRPV1-lineage processes presynaptic to Vc-parabrachial projection cells, whether mono- or polysynaptic connections link TRPV1 afferents to these neurons remains unclear. Nonetheless, neural messages from TRPV1-lineage fibers were selectively routed by the Vc to PB taste cells. Photoexcitation of TRPV1 afferents engaged only a minority of sweet neurons tuned to preferred tastes but stimulated most sodium, electrolyte, and bitter taste cells.

Excitation of TRPV1-lineage primary neurons represents a broad stimulus for somatosensory pathways that is functionally significant. TRPV1-lineage afferents comprise small diameter Aδ- and C-fibers that express peptidergic and nonpeptidergic nociceptive markers and project to superficial lamina in the dorsal horn of the spinal cord and Vc (Cavanaugh et al., 2011a; Mishra et al., 2011; Browne et al., 2017; Black et al., 2020), which relay to the PB area (Jasmin et al., 1997). Silencing TRPV1-lineage afferents in mice causes deficits in thermosensory, algesic-related, and pruritoceptive responses while excitation induces nocifensive behaviors (Mishra et al., 2011; Park et al., 2015; Browne et al., 2017). Behavioral assays of these fibers have focused on their roles in extra-trigeminal pathways. Nonetheless, trigeminal TRPV1-lineage afferents are implicated in orofacial pain-related responses in mice (Wang et al., 2017). The present results imply the functions of TRPV1-lineage fibers coursing the trigeminal tract are partly mediated by taste-active neurons in the brain.

### An organized overlap of taste and somatosensory processing in parabrachial circuits

Responses in PB neurons to TRPV1 afferent input, temperatures, and tastes were ordered in specific ways. While only sparingly engaging sweet neurons, photostimulation of TRPV1-lineage fibers frequently excited PB cells oriented to tastants capable of inducing aversion. Accordingly, responses to noxious hot temperatures emerged in PB heat neurons co-responsive to avoided bitter taste stimuli, similar to our recent results (Li and Lemon, 2019). In contrast, preferred sucrose and umami tastes strongly and selectively stimulated sweet neurons that showed some of the weakest responses to noxious heat. These features drove a unique positive association between PB population responses to bitter taste, noxious heat, and chemonociceptive stimuli of aligned aversive valence. Notably, these associations were lost in the simulated absence of TRPV1-lineage positive PB cells.

PB population responses to aversive bitter taste and nociceptive stimuli decorrelated with activity to moderate (10 mM citric acid and 100 mM NaCl) or preferred (sucrose and umami) tastes and with responses to oral cooling to 14°C. In brief-access tests, rats prefer to lick water chilled to 10°C over warm water (Torregrossa et al., 2012; Kay et al., 2020), implying some degree of oral cooling is not aversive to rodents. Further, wild-type mice show only a mild reduction from water in brief-access licking to the cooling agent menthol at the 1.28 mM concentration tested on PB neurons (Lemon et al., 2019), revealing mice do not avoid this stimulus. These data agree with the separation of menthol and cool temperatures from avoided bitter taste and nociceptive stimuli by PB responses.

Heat neurons equivalently responsive to noxious hot temperatures and bitter tastes represented the main intersection of thermosensory and gustatory sensitivity across PB cells (Figure 11). Importantly, neurons were selected for recordings based on sensitivity to taste solutions, but not temperature. Noxious heat may also engage non-gustatory PB neurons. Nonetheless, neural sensitivity to heat overlapped with responsiveness to bitter tastes, while innocuous cooling strongly stimulated PB cold cells with limited taste sensitivity. Notably, ordering neurons by sensory tuning revealed that PB neural clusters systematically differed from heat cells to reflect the valence of their most effective stimuli. Bitter neurons were most similar to and nearby heat cells while sweet neurons were furthest apart (Figure 11). Sodium, electrolyte, and cold neurons generally showed comparably intermediate divergence from heat cells. These trends associate with a neural code that could variably represent hedonic information for tastes and temperatures along a shared dimension.

**Figure 11.**
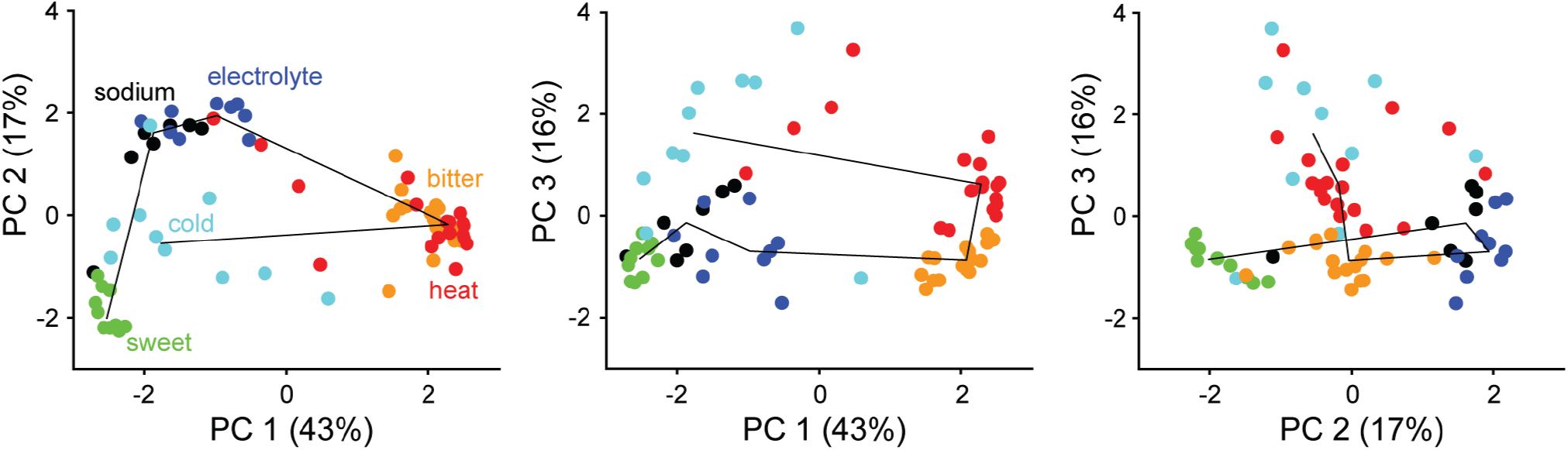
The transition of gustatory and thermal tuning across PB neurons captures cross-sensory valence. Principal component (PC) analysis visualized relationships between PB neurons (circles, *n =* 70) based on their responses to taste and temperature stimuli. Labels/colors reflect cell clusters in Figure 8B. Line connects the center points (median coordinates) of each cell cluster across dimensions. Parenthetical term gives variance explained by each PC. The arrangement of neurons reflects commonalties and differences in sensory tuning and the hedonic values of their most effective stimuli. Heat neurons co-responsive to aversive hot temperatures and bitter tastes are the main cellular intersection of gustatory and thermal sensitivity in PB circuits. Inspection of the first two PCs (leftmost plot), which capture the most variance, shows bitter neurons are the closest to heat neurons (most similar) with sweet neurons oriented to preferred tastes furthest apart (most dissimilar). Other cell clusters show comparably intermediate divergence.

We suggest the overlap of taste and temperature processing in PB circuits reflects a goal for parabrachial coding to capture aversive oral sensations regardless of modality or afferent pathway of signal arrival. Our study reveals that neurons traditionally defined as “taste cells” are part of this integrative process. Only when probed for sensitivity to somatosensory stimuli and excitation of somatosensory fibers was the broader response repertoire of these cells revealed. Notably, classic (Bernard and Besson, 1990; Bernard et al., 1994) and modern (Campos et al., 2018) studies show nociceptive and avoidance signals from diverse systems and body regions converge onto cells in the lateral PB area, where we targeted our recordings (Figure 1H). These and other results have kindled interest in the PB area as a center for protective processing and integration (Gauriau and Bernard, 2002; Palmiter, 2018; Chiang et al., 2019). It is curious if taste neurons in lateral PB regions respond to a larger array of cross-modal stimuli than presently tested.

Optically engaging TRPV1-lineage fibers revealed wider ties between functionally defined afferent input and PB activity than captured using external stimuli. While heat excited TRPV1-lineage positive PB neurons, few heat-responsive cells fired to lingual delivery of capsaicin. Capsaicin engages heat activated TRPV1 on primary nociceptive fibers (Caterina et al., 1997; Caterina et al., 2000). Sparse activity to capsaicin may be due its application to the rostral tongue. In contrast, heat and laser stimulation broadly stimulated, respectively, the oral mucosa and TRPV1 fibers within the trigeminal tract. Furthermore, because Cre recombinase is engaged in the embryo, ChR2 arose in the lineage of cells that express TRPV1 or transiently did so during development (Cavanaugh et al., 2011a; Mishra et al., 2011). Thus, non-TRPV1 mechanisms on TRPV1-lineage fibers may contribute to heat responses in PB gustatory cells.

TRPV1-lineage fibers communicated with diverse gustatory neurons, albeit noxious heat predominantly stimulated bitter taste-oriented cells. This may reflect input from a subpopulation of the nociceptive afferent types that comprise the TRPV1 lineage (Cavanaugh et al., 2011a). TRPV1-lineage input to electrolyte and sodium neurons may have roles unexamined here.

The axons of some TRPV1-positive trigeminal ganglion neurons bifurcate and project to both the Vc and to the PB area (Rodriguez et al., 2017). The possibility exists that photoexcitation of TRPV1-lineage terminals in the Vc could antidromically invade bifurcating trigeminal ganglion cells. Nevertheless, we used light power (∼5 mW at fiber tip) reported to not reliably induce antidromic action potentials in ChR2-positive axons (Jayaprakash et al., 2016).

### Parabrachial cold neurons can masquerade as broadly tuned taste cells

Our recordings encountered cold neurons that gave the strongest responses to oral cooling to 14°C. These cells showed limited taste sensitivity when responses were corrected for activity to the cooling ramp that accompanied taste delivery. However, cold cells significantly increased their breadth of responsiveness across taste solutions when response rates were uncorrected for cooling. Under this condition, cold neurons appear as “broadly tuned taste neurons” responsive to diverse tastes, although this effects was due to their thermal sensitivity. Nevertheless, significant effects of cooling on taste responses were observed only in cold neurons. While oral cooling frequently stimulated TRPV1-lineage positive PB cells, cold-evoked response rates were comparably small outside of cold cells.

### Looking ahead

The discovery that taste and somatosensation are represented together in the PB complex implies these modalities are only components of a larger neural code and there are dependencies between them. Future studies on the behavioral significance of gustatory and TRPV1 integrative neurons will be aided by identifying their genetic signature. Candidate neurons include cells that express calcitonin gene-related peptide, which mediate protective functions and occupy lateral PB regions (Campos et al., 2018; Palmiter, 2018) populated by TRPV1-lineage positive taste neurons. Delineating gustatory and TRPV1 integrative circuits will open new avenues of study on taste and sensory-integrative coding and may shed deeper light on pain-related processing in the PB nucleus.

## Acknowledgements

Supported by NIH grant DC 011579 to C.H.L. We thank Dr. Tingting Gu for assistance with microscopy.

